# Computational elucidation of possible contributors to formation and stabilization of ATP-lid down-conformation in the N-terminal domain of Hsp90

**DOI:** 10.1101/2025.02.27.640699

**Authors:** Keigo Gohda

**Affiliations:** Computer-aided Molecular Modeling Research Center, Kansai (CAMM-Kansai) Nishinomiya, Japan

**Keywords:** ATP lid, Hsp90, lid closure, molecular dynamic simulation, repulsive distance-restraint simulation

## Abstract

Heat shock protein 90 (Hsp90) controls activation and maturation of various crucial client proteins through a catalytic cycle. In this catalytic cycle, closure of the lid segment from up- to down-conformation in the N-terminal domain (NTD) of Hsp90 through ATP binding is indispensable for coordinated structural changes, including interchange of dimeric Hsp90 structure between open and closed forms. However, the mechanisms underlying lid closure remain unclear. In this study, we investigate structural characteristics of the lid-down conformation in an isolated monomeric NTD structure by two types of molecular-dynamic simulation: a flopping-down simulation for a lid up-conformation using repulsive distance-restraints, and a down-conformation simulation for in-silico H1-mutants of NTD with a lid-down conformation. In the flopping-down simulation, spontaneous formation of a lid-down conformation is observed in multiple times. K98 and K102 in the lid segment are observed to interact with ATP phosphate or D40, suggesting to contribute to formation of the lid-down conformation. In the down-conformation simulation, H1 structure of the chimera H1-model, which only retains a proper down-conformation among the models for the entire simulation period, covers over the lid segment more than that of the X-ray structure. Because stability of the lid-down conformation was influenced by H1 structures, H1 segment is suggested to contribute to stabilization of the lid-down conformation. Although no direct experimental data are currently available to confirm these findings, these simulation results do not show large discrepancies with the experimental data and evidence of structural characteristics of the NTD, deduced from previous X-ray and spectroscopic studies.

## 1. INTRODUCTION

The 90 kDa heat shock protein (Hsp90) is a chaperone protein that regulates the activation and maturation of various client proteins. These proteins play central roles in essential cellular processes involved in cell growth and proliferation through a catalytic cycle that hydrolyzes ATP molecules.^1–5^ Hsp90 functions as a homodimer and is composed of an N-terminal domain (NTD), including the ATP-binding site, a middle domain that mainly contributes to interactions with clients and co-chaperones, and a C-terminal domain (CTD), which is primarily involved in dimerization.^6–10^

In an apo state, that is, a state of no ATP molecules bound to the N-terminal ATP-binding site, the dimeric Hsp90 subunits adopt an “open form” only with the CTD dimerization.^11^ Binding an ATP molecule to the NTD drives conformational changes of the dimeric Hsp90 structure to a “closed form” with an NTD dimerization.^6^ In the closed structure, the NTD is bound to the middle domain and a catalytically important Arg330 located in the middle domain is directed to an ATP molecule in the NTD. The closed form activates the catalytic site, leading to ATP hydrolysis and the release of ADP molecules and mono phosphates from the dimeric subunits, therefore completing the catalytic cycle. One of the rate-limiting steps in the cycle involves coordinated NTD structural changes, including closure of the lid segment (Gly94-Gly123) in the ATP-binding site during the opening–closing process (yeast Hsp82 numbering is used throughout this manuscript). The closed lid segment covers over a bound ATP molecule at the ATP-binding site, as shown in **Figures 1a and b**.

**Figure 1.**
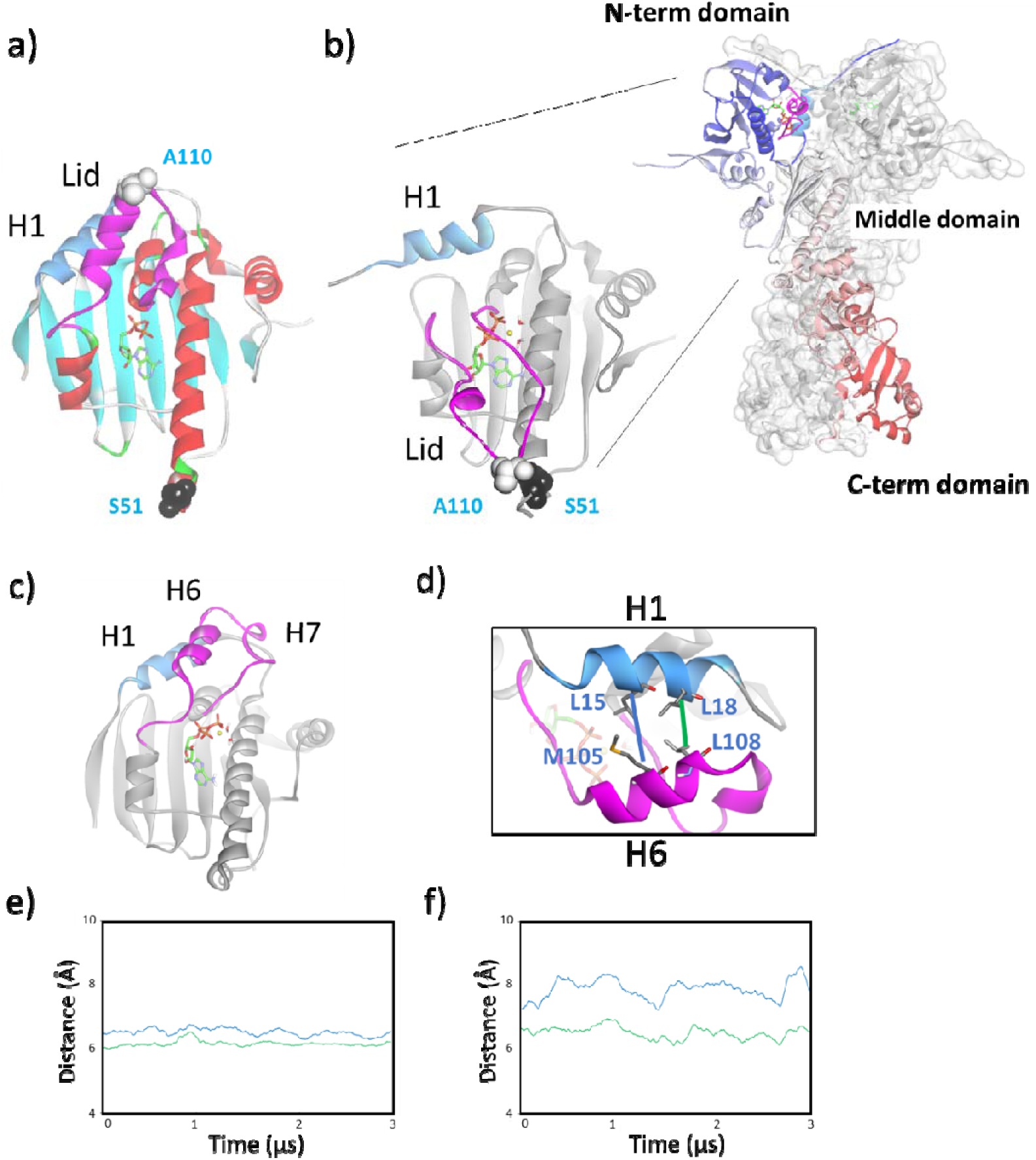
X-ray structures of yeast Hsp82 (Hsp90). a) Isolated NTD complexed with ADP (PDB ID: 1AM1). Lid (Gly94-Gly123) and H1 (Ala10-Asn21) segments are colored magenta and blue, respectively. b) Dimeric Hsp90 structure on the right and close-up view of the NTD in the left (PDB ID: 2CG9). Sba1 molecules in the complex are removed for clarity. Ser51 and Ala110 located at the apical end of the lid segment and body of the NTD are shown with black and white vdW models, respectively. These residues were used as chromophore probes in fluorescence resonance energy transfer (FRET) study.^14,34^ c) “15-μs up-conformation structure” obtained in our previous simulation study.^13^ d) Close-up view of H1–H6 segments. Distance trajectories of L15-M105 (blue lines) and L18-L108 (green lines) for e) wild-type and f) L15N/L18N structures.

As observed in X-ray and cryo-electron microscopic structures of monomeric or dimeric Hsp90 subunits, the lid segment exhibits an open conformation in the open form of the dimeric Hsp90 subunit and a closed conformation in the closed dimeric Hsp90 subunit, respectively.^1,11^ In this manuscript, the open and closed conformations of the lid segment are referred to as “up” and “down” conformations to avoid confusion with structural states of the dimeric Hsp90 subunit. Because the closure of the lid segment through ATP binding triggers ATP hydrolysis, this is a significant process in the catalytic cycle. The up- and down-conformational transition of the lid segment was recently reported in the NTD of human Hsp90α in the apo state through nuclear magnetic resonance (NMR) measurements.^12^ Although the lid up- and down-conformations have been clarified in the X-ray and cryo-electronic microscopic structures, the atomistic mechanism of the closure process of the lid segment has not yet been elucidated in an NTD complex with ATP.

To understand the closure mechanism, we previously investigated the structural transition from up- to down-conformation using 15-μs-length conventional molecular dynamic (MD) simulations for apo and ATP-complex structures of an isolated monomeric NTD of yeast Hsp90.^13^ Despite the long MD simulation, no lid closures were observed; however, a very early event, that is, *a sign* of lid closure leaving the core NTD structure was captured. By comparing the results of the same simulation for Hsp90 mutants, A107N and T101I, and their available experimental data (ATP hydrolysis activity and lid-closure experiment by ATP spiking using the fluorescence resonance energy transfer [FRET] method),^8,14^ it was concluded that lid closure from the up- to down-conformation is triggered by lid leaving accompanying the conformational instability of the lid segment such as helix-7 (H7) unwinding and slight displacement of the lid segment (**Figure 1c**).

As a subsequent study following our previous study, we further investigated the structural characteristic of the lid segment by focusing on the down-conformation. As described above, in X-ray and cryo-electron microscopic structures, a down-conformation has been observed in a dimeric Hsp90. However, an NMR study reported an existence of a down-conformation in a minor population, 3.2%, in an isolated monomeric human Hsp90.^12^ Because the apo structure of the NTD was used in this study, the lid-down conformation under an isolated monomeric NTD-ATP complex is not well characterized. In the present study, therefore, the characteristic of the lid-down conformation was computationally investigated using an isolated monomeric NTD structure to avoid any effect of other domains or molecules. The research questions addressed in this study are as follows: What is the key event or effect, if any, to achieve lid flopping from the up- to down-conformation and to stabilize the down-conformation in an isolated monomeric NTD structure?

## 2. MATERIAL and METHODS

### 2.1. Material

The last structure in the 15-μs simulation of the X-ray isolated yeast Hsp82 NTD structure was used as an initial structure of an isolated monomeric NTD with a lid up-conformation. This structure was obtained from in our previous study (referred to as the “15-μs up-conformation structure”).^13^ The structural preparation, including for the 15-μs simulation, is briefly described below. As the original X-ray structure is a complex with an ADP molecule (PDB ID: 1AM1),^15^ the positions of γ-phosphate, a magnesium ion, and two coordinating water molecules for the ATP complex were modeled from the human Hsp90α X-ray NTD structure complexed with ATP (PDB ID: 3T0Z).^16^ Structural information from previous studies on the conformation of catalytic residues was used for model building.^11,17^ The details of the model preparation are described in a previous report.^13^ For the initial structure of an isolated monomeric NTD with a lid-down conformation, the X-ray structure of yeast dimeric Hsp82-Sba1 complex was used (PDB ID: 2CG9).^6^ To prepare the structures for the MD simulation, the Sba1 molecule, water, and ions were removed, and one NTD structure was extracted from the X-ray dimeric structure. The positions of a magnesium ion and two coordinating water molecules were the same as used for the 15-μs up-conformation structure above. Hydrogen atoms were then added. The protonation states of the side-chain atoms were assessed using the PDB2PQR server.^18^ When assigning the protonation states, a neutral pH (7.0) was assumed. The in silico mutant/truncated structures were modeled by altering or deleting the amino acids at relevant positions in the X-ray structures. The model structures were manipulated using the MAESTRO modeling package (Schrodinger LLC, New York, NY, 2020). All molecular mechanics calculations were conducted using the *pmemd.cuda* or *sander* modules of AMBER24/AMBERTOOL24.^19–22^

For computations using AMBER24, the *tleap* module was used to prepare the topology and coordinate files of the proteins and ligands. The ff14SB force field and TIP3P model^23,24^ were used for protein and water molecules, respectively. The topology and parameter files of the ATP molecule and a Mg^2+^ ion were obtained from the AMBER parameter database of Bryce et al. at the University of Manchester.^25–27^ The modeled structures prepared using the *tleap* module were immersed in a cuboid box containing water molecules to impose a periodic boundary. The largest distance between the complex and water molecules in the box was set at 12.0 Å. To neutralize the entire system, Na^+^ counterions were added to the box.^28^

### 2.2. Methods

#### 2.2.1. MD Simulation

The modeled structure in the explicit solvent was subjected to energy minimization. The minimization procedure was divided into the following three steps with or without constraints: (a) The positions of all heavy atoms in the protein and ATP were fixed (positionally constrained with a 50 kcal/mol·Å^2^ weight), and hydrogen atoms, ionic atoms, and water molecules were allowed to move freely; (b) weak constraints were added to all heavy atoms in the protein and ATP (positionally constrained with a 2 kcal/mol·Å^2^ weight); and (c) no constraints were added. Each minimization process included 500 cycles of the steepest-descent method followed by the conjugate-gradient method. The energy convergence criterion gradient in each minimization step was 0.0001 kcal/mol·Å for the root mean square (RMS) of the Cartesian elements of the gradient. The cutoff distance for nonbonding interactions was 8.0 Å, and the nonbonding interaction list was updated every 25 steps.

The molecular dynamics calculations for the minimized structure were conducted in three consecutive steps: heating, density, and simulation. The time step in the MD simulation was 2 fs and the SHAKE constraint was used for bonds involving hydrogen atoms.^29^ The particle-mesh Ewald method was used to calculate electrostatic energy.^30,31^ The system was heated from 0 to 300 K over 100 ps using an Andersen temperature-coupling system.^32^ The velocities were randomized every 2 ps for temperature scaling. The Cα atoms of the main-chain atoms in a protein were positionally constrained by a 2 kcal/mol·Å^2^ weight. After heating, the system was equilibrated at 300 K at 1 bar for 50 ps. A constant-pressure periodic boundary condition was applied using a Berendsen barostat with a 1 ps pressure relaxation.^33^ The same positional constraints were applied to the system as in the heating step. Finally, a 50 ns simulation without any constraints was performed, and the obtained structures were applied for further MD simulations.

#### 2.2.2. Repulsive distance-restraint simulation

To invoke the flopping down of the lid segment, four inter-residual Cα-Cα distances were repulsively restrained: L15-M105, L18-L108, L18-M105, and L15-L108. Initial inter-residual distances were based on the “15-μs up-conformation structure”: Dist_L15-M105_=7.99 Å, Dist_L18-L108_=7.12 Å, Dist_L18-M105_=8.73 Å, and Dist_L15-L108_=9.19 Å.^13^

A distance of 1 Å was added to each of the four initial inter-residual distances as the current repulsive distance restraints, and a 1-μs simulation was conducted as the first cycle of the repulsive distance-restraint simulation. After the first simulation cycle, four inter residual distances were measured. Based on the results of the distance measurement, a new cycle of the simulation was implemented with either of the following three conditions: i) If all the measured inter-residual distances exceeded the current repulsive distance restraints plus an extra 1-Å distance, the repulsive distance-restraints were removed to zero, that is, no restraints for the new cycle. This situation was defined to be that a “free movement of the lid segment” has been emerged. ii) If at least one of the measured distances did not exceed the current repulsive restraints plus an extra 1 Å distance, 1 Å was added to all the current repulsive restraints as new distance restraints for the new cycle. iii) If at least one of the measured distances does not exceed the current repulsive restraints, the same current repulsive restraints were retained for the new cycle. For a post zero restraints cycle, the previous restraints before the zero-restraints cycle were used as the “current” restraints of the three conditions i)–iii). A 1-μs simulation was conducted for each cycle.

The cycle of the repulsive distance-restraint simulation was continued until the formation of a lid-down conformation or a structural change in the NTD, except for the lid segment, was observed. A “formation of lid-down conformation” was defined as an inter-residual Cα-Cα distance between Ser51 and Ala110 getting closer than the inter-residual distance observed in the X-ray NTD structure of down-conformation (PDB ID: 2CG9) (11.03 Å) plus 10 Å, that is, 21.03 Å (**Figure 1b**). Ser51 and Ala110, located at the apical ends of the lid segment and the body of the NTD, respectively, were used as residues, to which FRET chromophore probes were attached in the FRET study, to monitor lid flopping up and down in solution.^14^ The situation in which the formation of lid-down conformation in a time period when the free movement of the lid segment occurred was defined as an achievement of a “spontaneous formation of lid-down conformation.”

The first repulsive distance-restraint simulation, run #0, did not exactly follow this simulation protocol because this run was used to examine the length of a simulation cycle and the distance increment of the restraints (Section 3.1).

## 3. RESULTS

### 3.1. Flopping down of the lid segment in an isolated monomeric NTD-ATP complex using repulsive distance-restraint MD

We first investigated the process of changing the lid segment from the up-to-down conformation. In our previous study, a flopping down of the lid segment was not observed during the 15-μs length MD simulation starting from the X-ray isolated NTD structure with the up-conformation.^13^ By analyzing the last structure of the lid conformation in the 15-μs NTD simulation, that is, the “15-μs up-conformation structure,” it was shown that a flopping down of the lid segment needs the lid to leave from the H1 segment (Ala10-Asn21). In particular, two hydrophobic residues in each H1 segment and H6 of the lid segment are located at an interface between the H1 and H6 segments (L15 and L18 in H1, and M105 and L108 in H6) with close Cα-distances of L15-M105 (6.9 Å) and L18-L108 (5.8 Å) (**Figure 1d**). Because this hydrophobic interaction seemed to prevent the lid from leaving the H1 segment, an in silico NTD mutant, replacing the two hydrophobic residues at the H1 segment with hydrophilic Asn residues, L15N/L18N, was modeled to examine the stability of the hydrophobic interaction. The L15N/L18N mutant model and the wild-type 15-μs up-conformation structure were subjected to a 5-μs simulation (**Figures 1e and f**). As a result, the H1-H6 interaction of the H1 hydrophilic mutant fluctuated, that is, there were larger deviations in both distances, compared with the wild-type (**Table S1**), thus indicating that the interaction between the H1 and H6 segments is unstable in the mutant.

Based on this result, the hydrophobic interaction between the H1 and H6 segments was suggested to glue the lid segment to the up conformation. As such, it was assumed that the expansion of the H1-H6 distances leads to a conformational transition from up-to-down conformation. To expand the H1-H6 distance, a repulsive distance-restrained MD simulation was used as a flopping-down simulation of the lid segment. Four repulsive distance restraints between the hydrophobic residues at H1 and H6 segments were introduced to a simulation of the wild-type 15-μs up-conformation structure; four inter-residual Cα distances were on L15-M105, L18-L108, L15-L108, and L18-M105 (**Figure 2a**). During most of the cycles in the simulation, the repulsive distance restraints were increased stepwise by 1 Å per 1-μs simulation until a “formation of lid-down conformation,” which was defined as the FRET residual distance, Ser51-Ala110, getting closer than the distance in the X-ray down-conformation NTD structure plus 10 Å, or a structural change of the NTD, except for the lid segment, was observed (Section 2.2.2). The 15-μs up-conformation structure, but not the X-ray isolated NTD structure with the up-conformation, was used as the initial structure for the flopping-down simulation since the structure and interaction of H1-lid segment should be well equilibrated and stabilized. Initial distances were based on the inter-residual distances (7.12 to 9.19 Å) of the 15-μs up-conformation structure.

**Figure 2.**
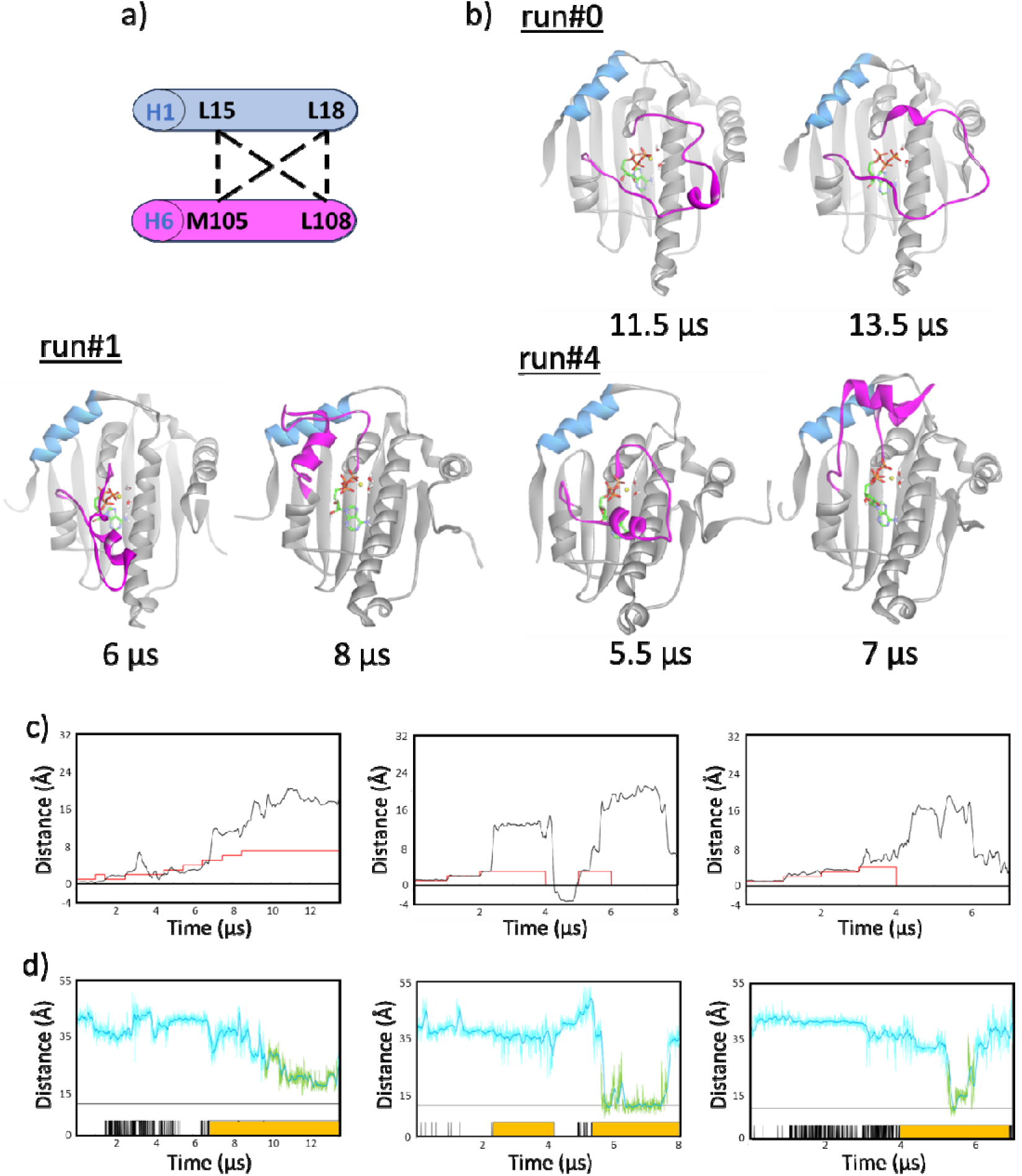
Successful flopping-down simulation of the NTD structure with up-conformation: runs #0, #1, and 4. a) Four repulsive distance restraints between H1 and H6 segments. b) Representative and the final structures of flopping-down simulations. c) Mean trajectories of the four restrained distances (black lines) and distance restraints (red lines). d) Trajectory of S51–A110 distance (light-blue lines). Trajectory shown in light-green is the “formation of the down-conformation” period (Section 2.2.2). The “free movement of the lid segment” periods are denoted with black vertical lines and the continuous periods are shown with orange bars (Section 2.2.2). c, d) Results of runs #0, #1, and #4 are shown on the left, middle, and right, respectively.

The repulsive restraint simulation was conducted in seven times (**Table 1**). The first run (run #0) was used to examine the length of the simulation cycle and the distance increments of the restraints in the simulation course. Of the seven runs, three runs, that is, runs #0, #1, and #4, achieved a “spontaneous formation of lid-down conformation” (**Figure 2b**). In run #0, a “free movement of the lid segment,” which was defined as the inter-residual distances exceeding the repulsive distance-restraint values plus 1 Å, was seen from around 7 μs at the cycle with 7-Å distance restraints as shown in **Figures 2c and d**. From around 9.7 μs to the end of the simulation (13.5 μs), the formation of the lid-down conformation was continued (**Figure 2d**). As this formation of the lid-down conformation occurred during the free movement period, a “spontaneous formation of lid-down conformation” was considered to have emerged, as was defined in the Material and Methods Section. The structure of the lid-down conformation was slightly expanded to a circular form compared to the X-ray down-conformation structure (**Figures 2b and S1a**). In run #1, the lid segment also adopted the down-conformation around 5.7 to 7.7 μs within the period of the free movement, that is, a spontaneous formation of lid-down conformation, and moved up thereafter (**Figures 2b–d and S1b**). The structure of the down-conformation was twisted, and the right side of the lid structure was lifted. In run #4, a rather short spontaneous formation of lid-down conformation was observed around 5.3 to 6.0 μs (**Figures 2b–d and S1c**).

**Table 1.** Results of H1-H6 repulsive distance-restraint simulation.

| **Runs** | **Formation of**<br>**down-conformation<sup>a)</sup>** | **Retention time of**<br>**down-conformation<sup>b)</sup> (μs)** | **Total**<br>**simulation**<br>**length (μs)** | **Maximum**<br>**repulsive**<br>**distance-restraints**<br>**($\text{\AA}$)** |
| --- | --- | --- | --- | --- |
| #0 | formed | 3.8 | 13.5 | 7 |
| #1 | formed | 2.0 | 8 | 3 |
| #2 | failed:<br>adenine-ring flipped | - | 6 | 4 |
| #3 | failed:<br>lid upward | - | 12 | 7 |
| #4 | formed | 0.7 | 7 | 4 |
| #5 | failed:<br>H1-length expanded | - | 16 | 9 |
| #6 | failed:<br>H1-position shifted | - | 24 | 17 |

a) Formation of down-conformation is defined as that the distance between Cα -atoms of Ser51 and Ala110 becomes closer than 21.03 $\text{\AA}$ .
b) Retention time was calculated by subtracting the start time of down-conformation formation from the end time.

The other four runs (run #2, #3, #5, and #6) failed to achieve lid-down conformation (**Table 1**). In the failed simulations, conformational aberrations in the NTD structure were observed, which led to the termination of the simulation, and no spontaneous formation of the lid-down conformation emerged (**Figure S2**). Run #2 showed flipping of the adenine ring in the ATP molecule at the end of the simulation (**Figures S2 and S3a**). The lid segment in run #3 moved upwards, probably because the distance restraint pushed the lid segment in the upper direction (**Figures S2 and S3b**). In runs #5 and #6, H1 structures were deformed at the end of simulations; the length of the H1 segment was extended along with absorbing an N-terminal β-strand structure in run #5, and the relative location of the H1 segment was shifted upward in run #6 (**Figures S2 and S3c and 3d**).

To find the possible interaction or movement required for the spontaneous formation of the lid-down conformation, the trajectories of the flopping-down simulation were visually inspected. In the successful simulations, the side chains of two lysine residues, K98 and K102, in the lid segment were observed to interact with the phosphate group of ATP or the carbonyl side chain of Glu40 in flopping-down conformations (**Figure 3**). The interaction of K98 with the ATP phosphate has been reported in the previous 15-μs simulation of the X-ray NTD structure with lid up-conformation, indicating that the K98-ATP interaction is used to hold the ATP phosphate to an appropriate position for the catalytic reaction (**Figure 3d**).^13^ A comparison of the distance trajectories of the two interactions, K98-ATP phosphate and K102-ATP phosphate, or D40 (as shown with the black and red lines in **Figures 3a–c**, respectively), with the FRET distance (as shown with the light-blue lines in **Figures 3a–c**) showed that the formation of the down-conformation (as shown with the light-green lines in **Figures 3a–c**) occurred when both the K98 and K102 interactions were simultaneously formed.

**Figure 3.**
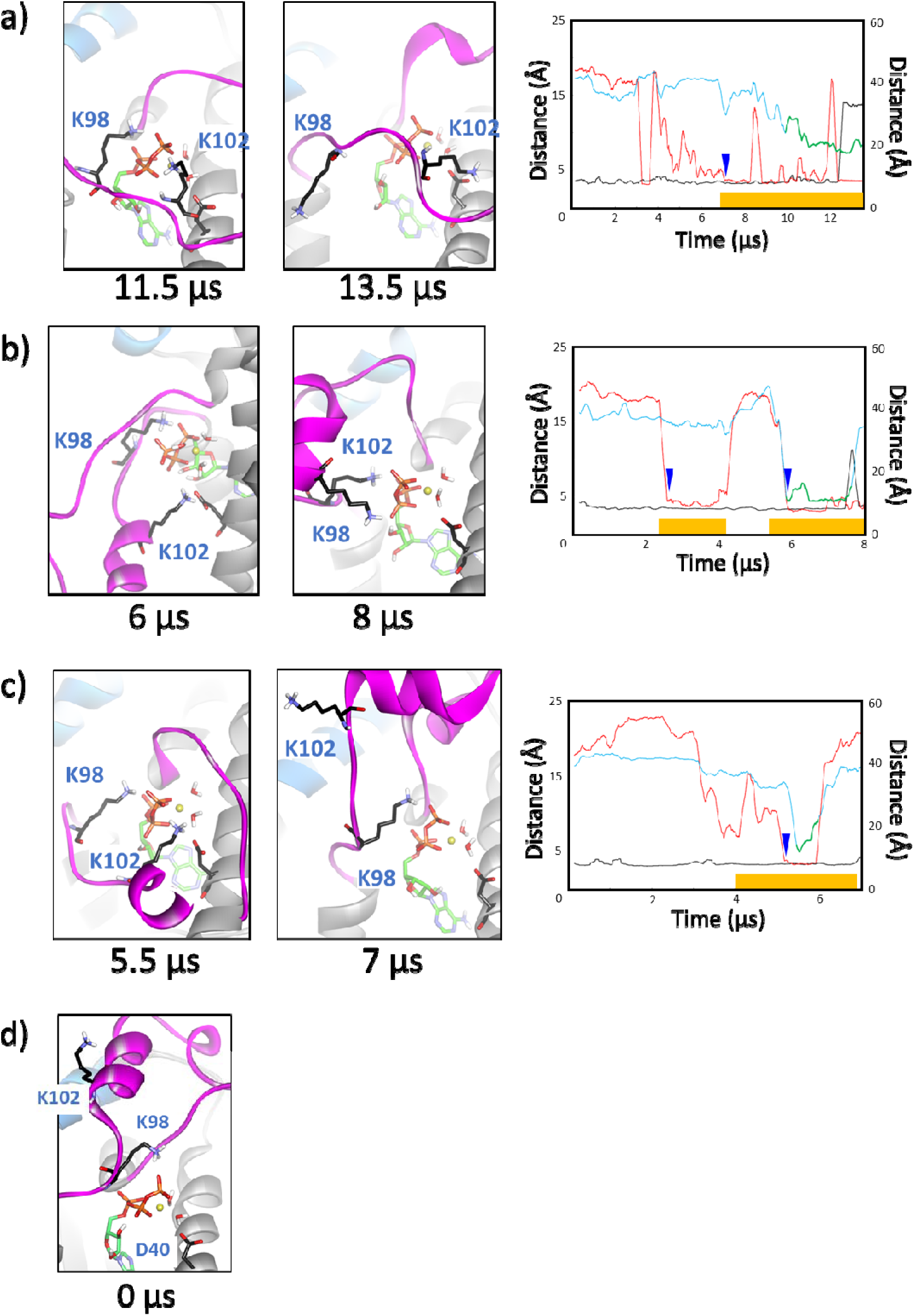
Close-up of representative and the final structures of the flopping-down simulation, a) runs #0, b) #1, and c) #4. d) Initial structure (0 μs) for each simulation (the 15-μs up-conformation NTD structure). a–c) Distance trajectories of K98-ATP phosphate (black lines) and K102-ATP phosphate/D40 (red lines) are shown in the left vertical axis. Distance trajectory of S51-A110 (light-blue lines) is shown on the right vertical axis. Light-green lines and orange bars indicate periods of “formation of the down-conformation” and “free movement of the lid segment,” respectively, also shown in Figure 2d. Blue wedges in the graphs indicate a starting point of the simultaneous formation of the K98 and K102 interactions under the free movement period of the lid segment.

In failed simulations showing no spontaneous formation of the down-conformation, the K98/K102 interaction was sporadically observed (**Figure S4**). Although in run #5 K98 and K102 interactions were simultaneously observed under the free movement period of the lid segment at 14–15 μs (**Figure S4d**) and the other runs had K98 and K102 interactions in some periods of time (**Figures S4b–c and S4e**), no flopping down of the lid segment occurred. This is probably because the free movement period was not sufficiently long to invoke the lid conformation in the simulations.

These flopping-down simulations indicate that K98/K102 is associated with the closure of the lid segment.

### 3.2. Stability of the lid-down conformation in an isolated monomeric NTD-ATP complex

The lid-down conformation in an NTD-ATP complex is a key structure that induces a series of structural changes in the catalytic cycle, such as association of the middle domain with the NTD and formation of a dimeric NTD. The stability of the lid-down conformation in an isolated monomeric NTD was thus investigated; however, no experimental structures of the lid-down conformation were obtained in an isolated monomeric NTD. Therefore, an X-ray dimeric NTD structure with a lid-down conformation was used as an isolated NTD structure.

A monomeric NTD structure was extracted from the X-ray dimeric NTD structure (named as “X-ray H1-model”) and subjected thrice for a 5-μs simulation (**Figure 4**). As a result, run #1 retained a down-conformation until 5 μs, but in the other two runs, runs #2 and #3, the lid segment moved upward before 5 μs, resulting in termination of the simulation at 2 μs for run #2 and 1 μs for run #3, respectively. In run #1, although the lid segment was kept in a down-conformation, the lid structure was twisted, and the right side of the lid structure was lifted (**Figures 4 and S5**). In run #2, the lid segment moved upwards and adopted a protruding helical conformation (**Figures 4 and S6**).

**Figure 4.**
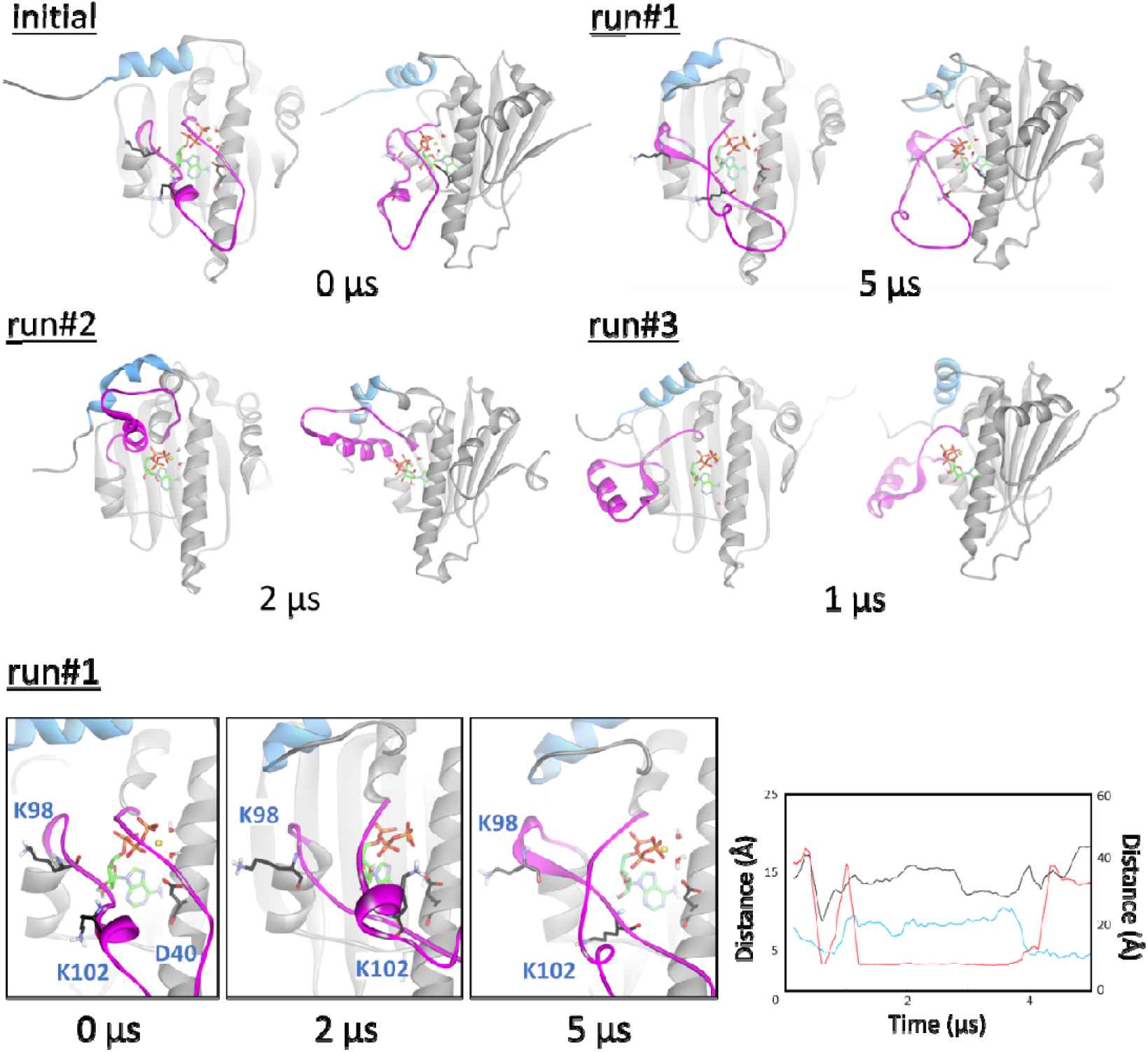
Initial and final structures of the down-conformation simulation, runs #1–#3, for the X-ray H1-model from the front and side. Close-up views of run #1 are shown at the bottom. In the graph, distance trajectories of K98-ATP phosphate (black lines) and K102-ATP phosphate/D40 (red lies) are shown on the left vertical axis. Distance trajectory of S51-A110 (light-blue lines) is shown on the right vertical axis.

This protruding conformation appeared to be blocked by the H1 segment, preventing further upward movement. In run #3, the lid segment swayed toward the side and formed a helical structure. The interactions of K98 and K102 with ATP phosphate or D40, which were suggested above to be associated with closure of the lid segment, were examined in the trajectories of the simulations. In the initial structure (at 0 μs) of the simulation, that is, the X-ray dimeric NTD with the lid-down conformation, no K98 and K102 interactions were observed (**Figure 4)**. In runs #1 and #2, K98 or K102 interaction was observed during most part of the simulations, but no such interactions were observed in run #3 (**Figures 4 and S6**).

To further understand the down-conformation stability, the final structures of the X-ray H1-model simulation were compared (**Figure 5a**). As a result, the position of the H1 segment varied among the three simulated structures, although no distance restraints that affected to the H1 structure in the flopping-down simulation were applied in this simulation. This indicates that the H1 structure influenced the lid conformation or vice versa during the simulation. To clarify the effects of the H1 structure on lid conformation, the following two H1 structures were modeled: “chimera” and “truncated” H1-models (**Figure 5b**). The chimera H1-model replaced the H1 structure of the X-ray H1-model with that of the 15-μs up-conformation structure to resemble an isolated monomeric NTD-ATP structure just after complexing with an ATP molecule. In the truncated H1-model, a whole H1 segment was removed to examine the intrinsic stability of the lid-down conformation. These modeled NTD structures were subjected to a 5-μs simulation.

**Figure 5.**
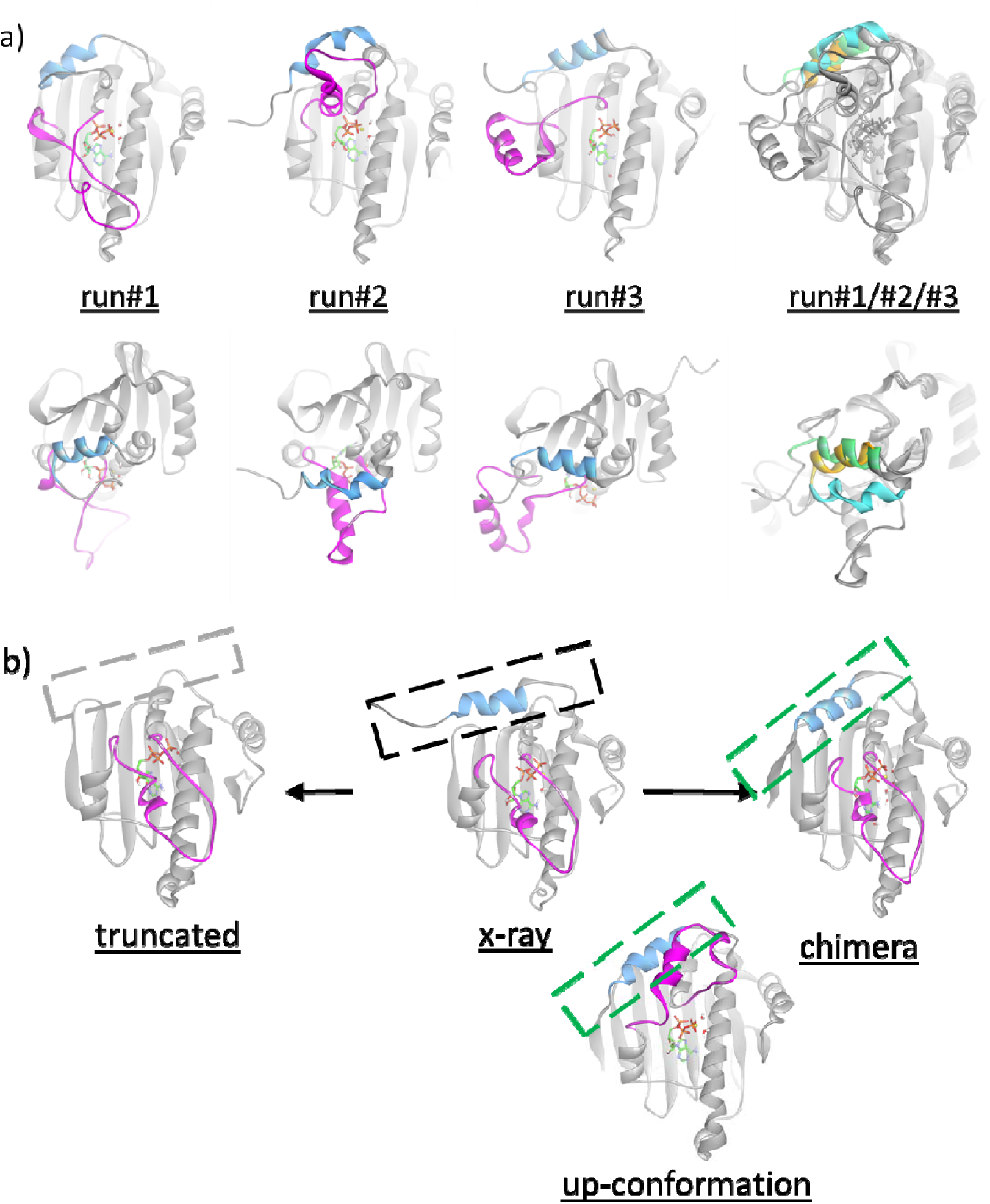
a) Comparison of the final structures of the down-conformation simulation, runs #1–#3, for the X-ray H1-model from the front (top) and side (bottom) views. At the right, the three final structures of runs #1–#3 are superimposed. H1 segments are colored light-yellow for run #1, light-blue for run #2 and light-green for run #3. b) Schematic image of H1-modeling: X-ray, chimera, and truncated H1-models.

The simulation of the chimera H1-model was duplicated and yielded similar results (**Figures 6 and S7**). In runs #1 and #2, the down-conformation of the lid segment was stably retained during the entire course of the simulation (**Figure S7**). Lifting of the lid segment was not observed, and K98 and K102 interactions were not observed in the simulations (**Figure 6**). Superimposing the initial and final structures of run #1 showed that the position of the H1 segment in run #1 moved down on the lid segment compared to that in the initial structure. In addition, the H1 position of run #1 was located between the two H1 segments of the dimeric NTD structure (**Figure S8**). For the truncated H1-model, a 5-μs simulation was conducted thrice, runs #1, #2 and #3 (**Figure 7**). Although the truncated structure had no H1 segment, the lid segment in run #1 retained its down-conformation until the end of the simulation, whereas the lid structure was twisted and the right side of the lid segment was lifted (**Figures 7 and S9**). For runs #2 and #3, the lid segment promptly moved upward within 1 μs, and then the simulations were terminated. As for the K98/K102 interaction, run #1 formed a K102 interaction, but not a K98 interaction. In run #2, K98/K102 interactions were partially observed, whereas no K98/K102 interactions were observed in run #3 (**Figures 7 and S9**).

**Figure 6.**
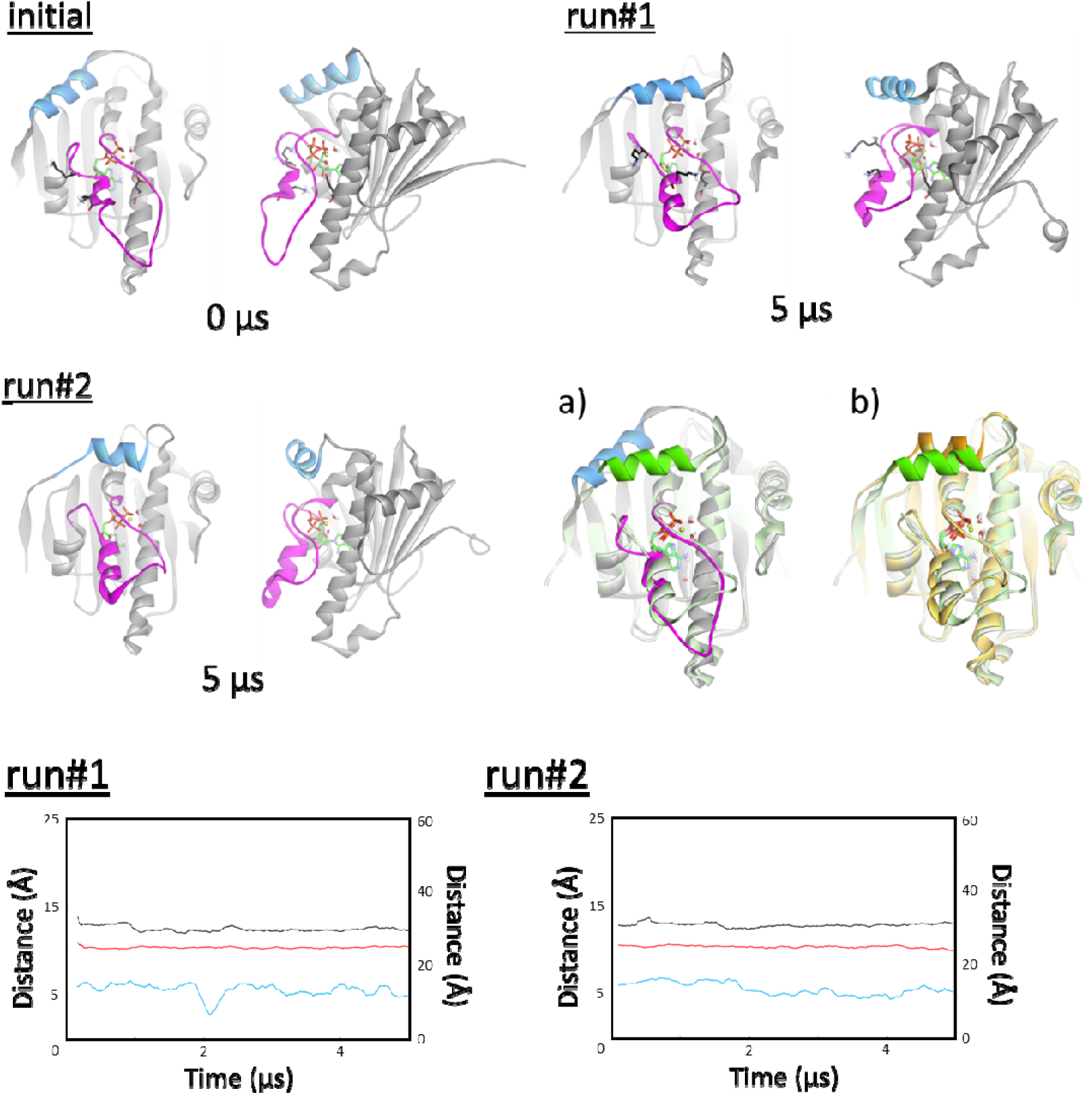
Initial and final structures of the down-conformation simulation, runs #1 and #2, for the chimera H1-model from the front and side. a) Superimposition of the initial and final structure of run #1 (green) and b) superimposition of the final structures of runs #1 (green) and #2 (orange). In the graphs, distance trajectories of K98-ATP phosphate (black lines) and K102-ATP phosphate/D40 (red lies) are shown on the left vertical axis. Distance trajectory of S51-A110 (light-blue lines) is shown on the right vertical axis.

**Figure 7.**
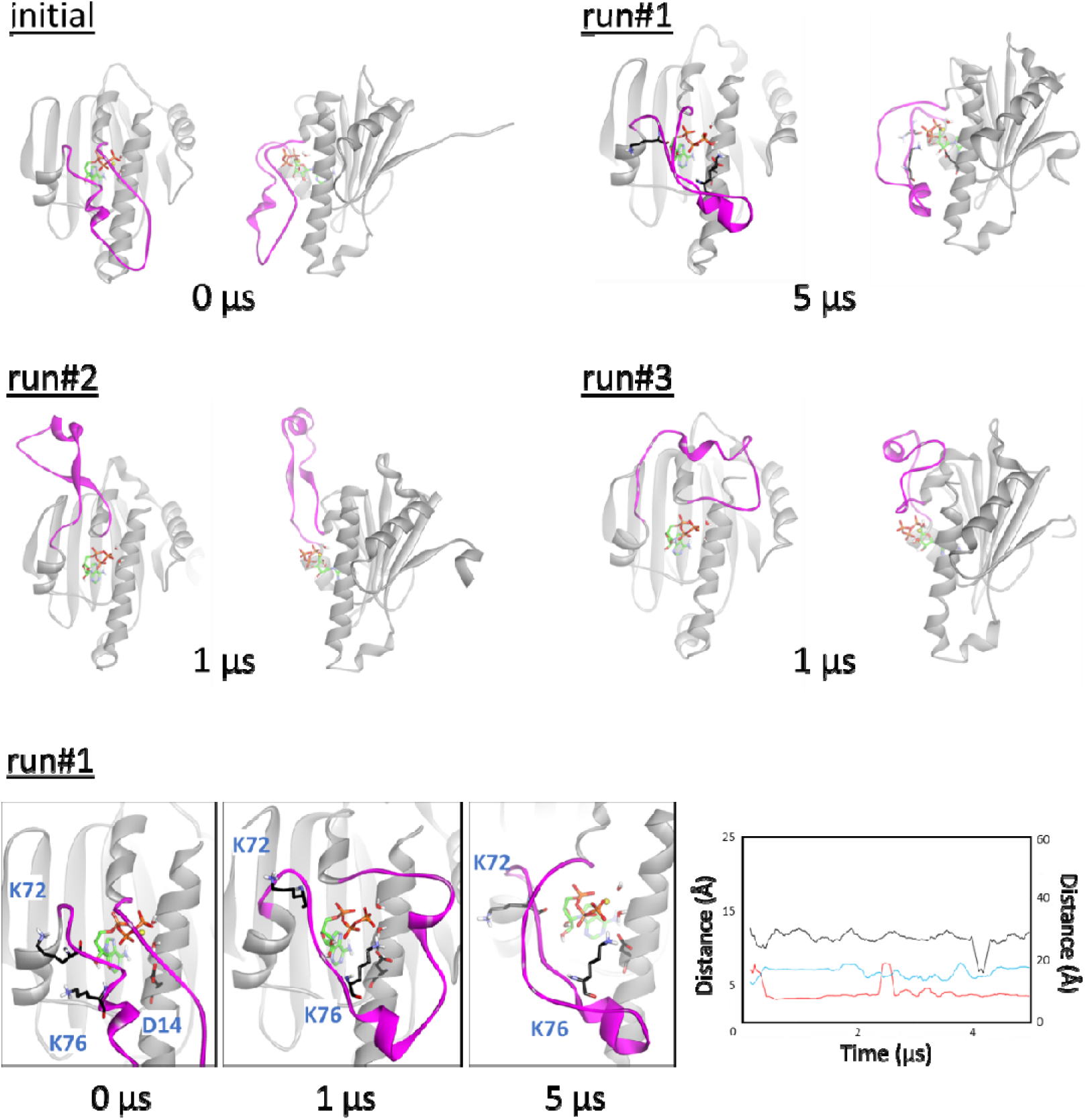
Initial and final structures of the down-conformation simulation, runs #1–#3, for the truncated H1-model from the front and side views. Close-up views of run#1 are shown at the bottom. In the graphs, distance trajectories of K98-ATP phosphate (black lines) and K102-ATP phosphate/D40 (red lies) are shown on the left vertical axis. Distance trajectory of S51-A110 (light-blue lines) is shown on the right vertical axis.

The H1-model simulations using the X-ray, chimera, and truncated models, indicates that the H1 structure contributes to the stability of the lid-down conformation in an isolated monomeric NTD structure, whereas the K98/K102 interaction may not.

## 4. DISCUSSION

Hsp90 controls the activation and maturation of various crucial client proteins through a catalytic cycle. The catalytic cycle involves molecular associations and structural changes in various modules and co-chaperone proteins. Among the molecular and structural events, ATP binding to the NTD triggers the implementation of the catalytic cycle, especially by inducing changes in the NTD, such as closure of the lid segment, association of the middle domain, and dimerization (swapping of the N-terminal segment) of the NTD.^14,34^ In a recent study, we addressed the mechanism of lid conformational changes to understand the initiation of the catalytic cycle at an atomic scale.^13,35^ Our previous study using 15-μs MD simulations on the lid transition from up- to down-conformation in an isolated monomeric NTD structure showed that an H7 unwinding and displacement of the lid segment may be a very early event of the lid closure, whereas no lid closures were observed in the long 15-μs simulation.^13^

In the present study, to understand the structural characteristics of the lid-down conformation, we conducted two types of MD simulations: a flopping-down simulation using repulsive distance restraints between the H1 and lid segments, and a down-conformation simulation using the H1-models. The former is to examine the formation of the down-conformation, and the latter is to assess the stability of the down-conformation. Both simulations apply for an isolated monomeric NTD structure in order that the investigations on the lid-down conformation are conducted without any effect of other domains or molecules. For the flopping-down simulation, a steered MD method, which has been widely used to study structural changes,^36^ was not applied. This is because a proper flopping-down pathway of the lid segment has not been elucidated, and spontaneous flopping down, that is, a motion without any external forces, is considered indispensable for exploring a mechanism for forming a down-conformation.

In the flopping-down simulation, after or at the same timing when both the K98 and K102 interactions with ATP phosphate/D40 were simultaneously formed under the state of the free movement of the lid segment, a flopping-down of the lid segment was commonly observed as long as the free-movement period was sufficiently long (blue wedges in **Figures 3 and S4**). As a direct interaction of K98 and K102 was observed, a possible association of K98/K102 was suggested for the formation of the lid-down conformation. A comparison of the corresponding residues to K98/K102/D40 in yeast and human Hsp90 homologues, whose structures are currently available, shows that the spatial positions of all residues are well agreed upon among the Hsp90 homologues: yeast Hsp82, yeast Hsc82, human Hsp90α, human Hsp90β, human TRAP1, and human Grp94 (**Figure S10**). This spatial conservation of the residues may support the roles of K98 and K102 in Hsp90. In addition, assuming that the ATP phosphate is involved in the K98/K102 interaction, ATP binding to the NTD directly facilitates the formation of the down-conformation, which agrees with experimental evidence that ATP binding triggers structural changes in the NTD structure.^14,34^

However, the K98/K102 association is not a major contributor to stabilizing the lid-down conformation in an isolated monomeric NTD because the down-conformations achieved in the flopping-down simulation (runs #1 and #4) moved upward at the end of the simulation. For the stability, it was indicated from the down-conformation simulation that the H1 structure may contribute because the stability of the lid-down conformation was influenced by H1 structures; the H1 segment of the final structure of the chimera H1-model, which covered over the lid segment more than that of the X-ray H1-model, only retained a proper down-conformation for the entire simulation period among the three different H1-models. In the X-ray dimeric Hsp90 structure, the lid-down conformation was sandwiched between the middle domain and the H2 segment of the NTD (**Figure S11**). To become sandwiched with the middle domain, the lid segment should first form a down-conformation and keep it in a certain period of time in the state of an isolated monomeric NTD-ATP complex until the middle domain comes into contact. Thus, the long-lasting down-conformation observed in the chimera H1-model agrees with this experimental observation.

An interesting observation in this study was the finding of a *partial* lid-down conformation, that is, a down-conformation with twisting and lifting of the right side of the lid segment. This structure was observed both in the flopping-down simulation (run #1) and in the down-conformation simulation (X-ray and truncated H1-models). In the partial lid-down conformation, the K102 interaction is commonly observed, whereas the K98/K102 interaction has not been observed in the *full* down-conformation of the dimeric Hsp90 structure or the chimera H1-model. The contribution of the K98/K102 interaction to the partial down-conformation is not sufficiently clear, but K102 may stabilize the partial down-conformation to a certain extent because the K102 interaction is observed even in the simplest H1-model, the truncated H1-model, in this study.

In conclusion, we investigated the structural characteristics of the down-conformation in an isolated monomeric NTD structure using various MD simulations and determined that K98/K102 and H1 segment are possible contributors to the formation and stabilization of the down-conformation, respectively. Although no direct experimental data are currently available to confirm these findings, the simulation results do not show large discrepancies with the experimental data or evidence of the structural characteristics of the NTD, deduced from previous X-ray and spectroscopic studies. However, further investigations, such as mutation and spectroscopic measurements, are required for clarifying these.

In future perspective, a small ATP-site binder interacting with K98/K102 may facilitate middle-domain association and/or NTD dimerization through the induction of a lid-down conformation, eventually leading the NTD to an active closed-form of Hsp90. This type of small molecule can then be an Hsp90 positive modulator like an activator co-chaperone and would be capable of being used for treatment of cancer or neurodegenerative diseases such as Alzheimer’s disease and Lewy body dementia.

## Disclosure statement

No potential conflict of interest exists.

## Funding information

The author reports no funding associated with the work featured in this article.

## Peer review

The peer review history for this article is available at https://publons.com/publon/xxxx.

## Data availability statement

The data that support the findings of this study are available from the corresponding author upon reasonable request.

## Abbreviations

FRET: fluorescence resonance energy transfer
Hsp90: 90 kDa heat shock protein
MD: molecular dynamics
NTD: N-terminal domain.

## SUPPORTING INFORMATION

Additional supporting information may be found online in the Supporting Information section at the end of this article.

## Notes

### Competing Interest Statement

The authors have declared no competing interest.

