## Supplementary material for "Computational elucidation of possible contributors to formation and stabilization of ATP-lid down-conformation in the N-terminal domain of Hsp90": Table S1, Figures S1 to S11

### TABLES

**Table S1.** Mean distances of H1–H6 segments in L15N/L18N mutant and wild-type structures in the 5-μs simulation.

| | **Mean distance (Å) ± SD ($Å^2$)** | |
|---|---|---|
| | **L/N15-M105** | **L/N18-L108** |
| L15N/L18N mutant | 7.88 ± 0.8506 | 6.55 ± 0.6441 |
| L15/L18 wild type | 6.54 ± 0.5067 | 6.17 ± 0.4130 |

SD = standard deviation.

### FIGURES and FIGURE LEGENDS

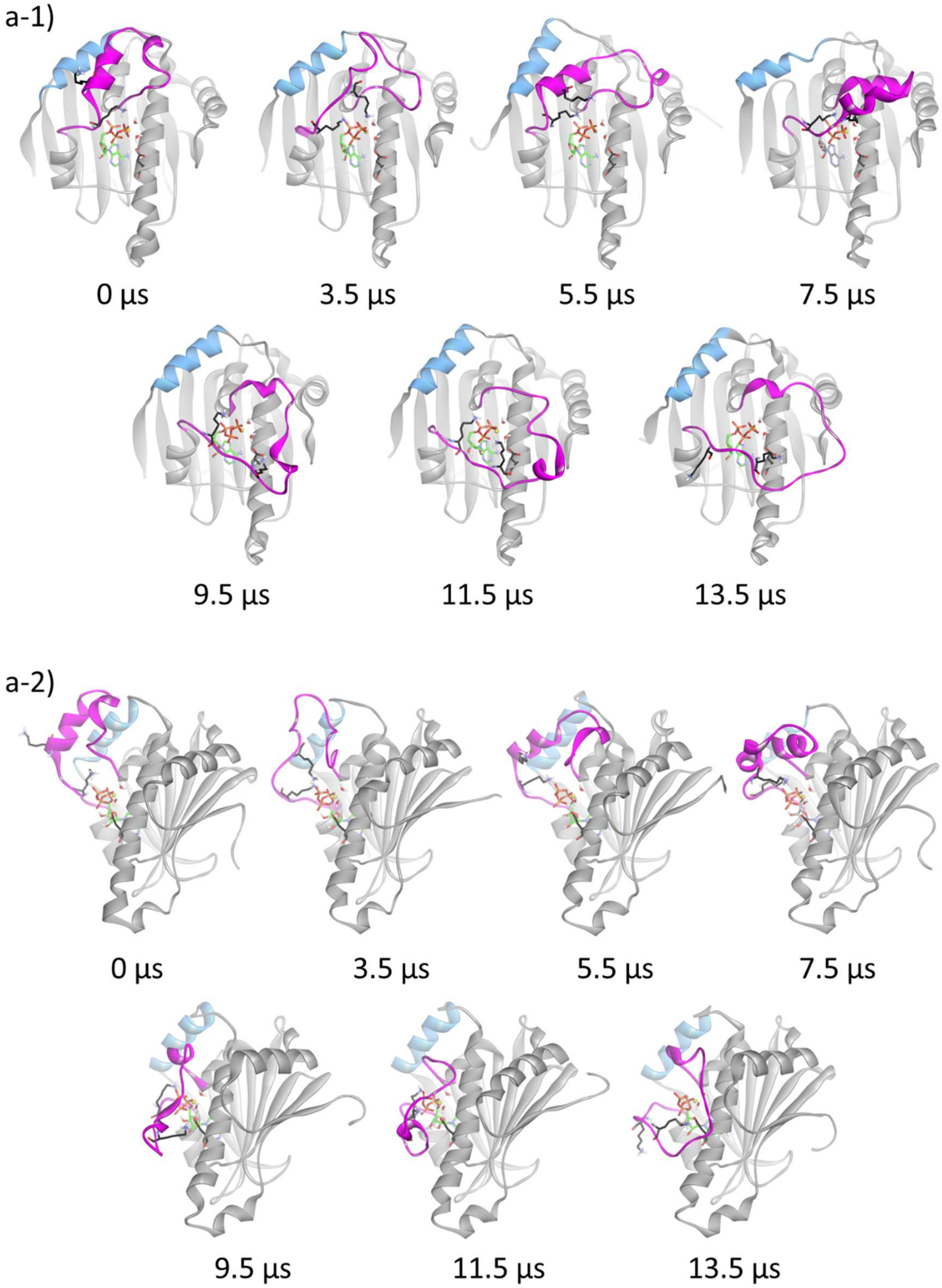


**Figure S1.** Trajectory snapshot structures of the flopping-down simulation, a) runs #0, b) #1, and c) #4 from the front (branch number: 1) and side (branch number: 2) views.

b-1)

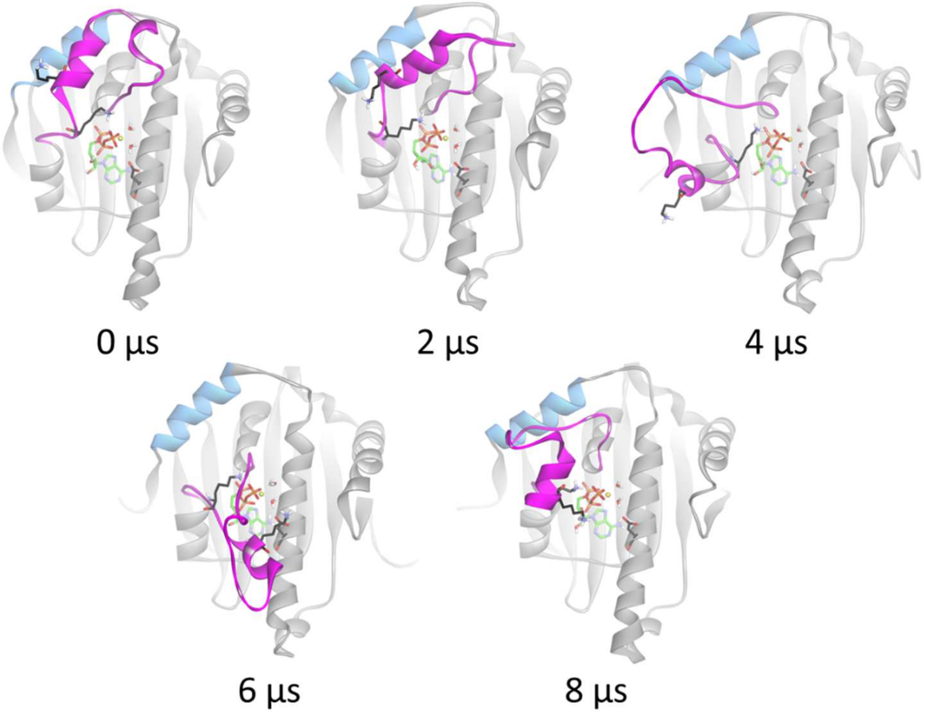


b-2)

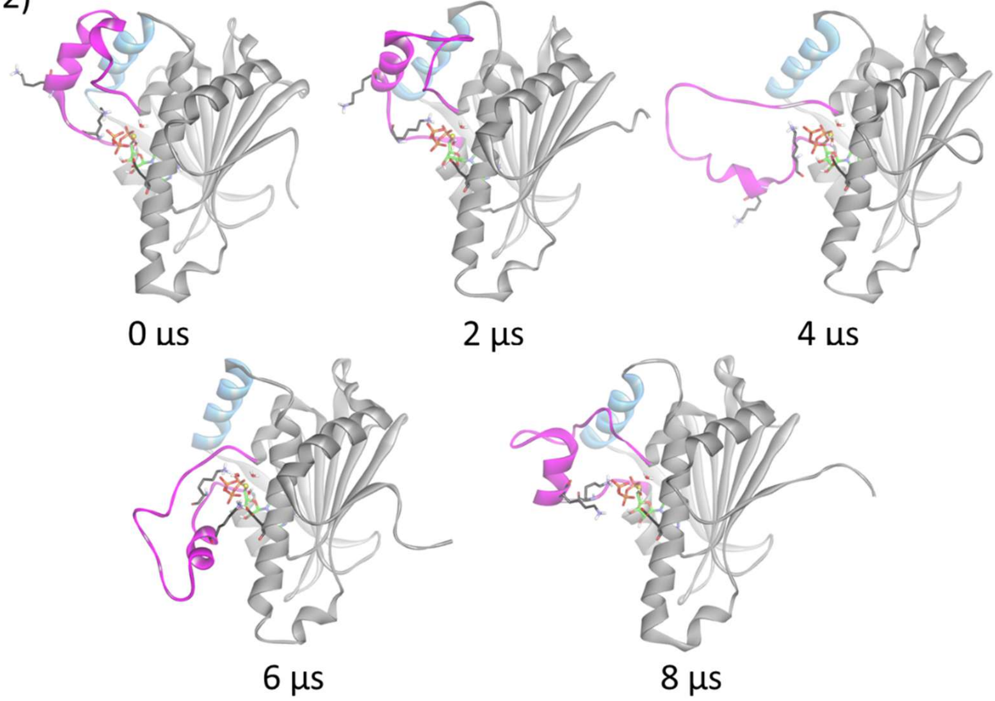


**Figure S1.** Trajectory snapshot structures of the flopping-down simulation, a) runs #0, b) #1, and c) #4 from the front (branch number: 1) and side (branch number: 2) views.

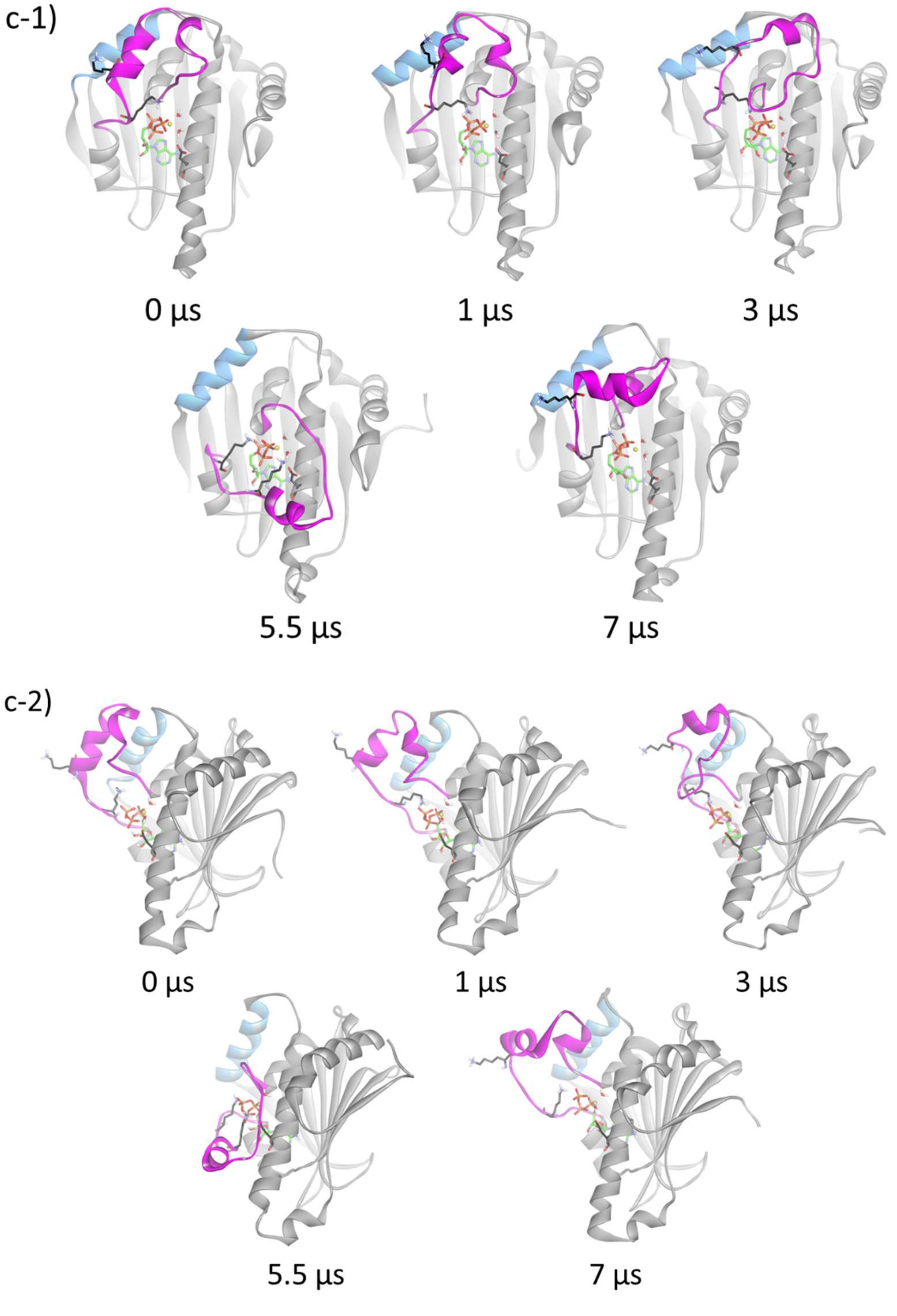


**Figure S1.** Trajectory snapshot structures of the flopping-down simulation, a) runs #0, b) #1, and c) #4 from the front (branch number: 1) and side (branch number: 2) views.

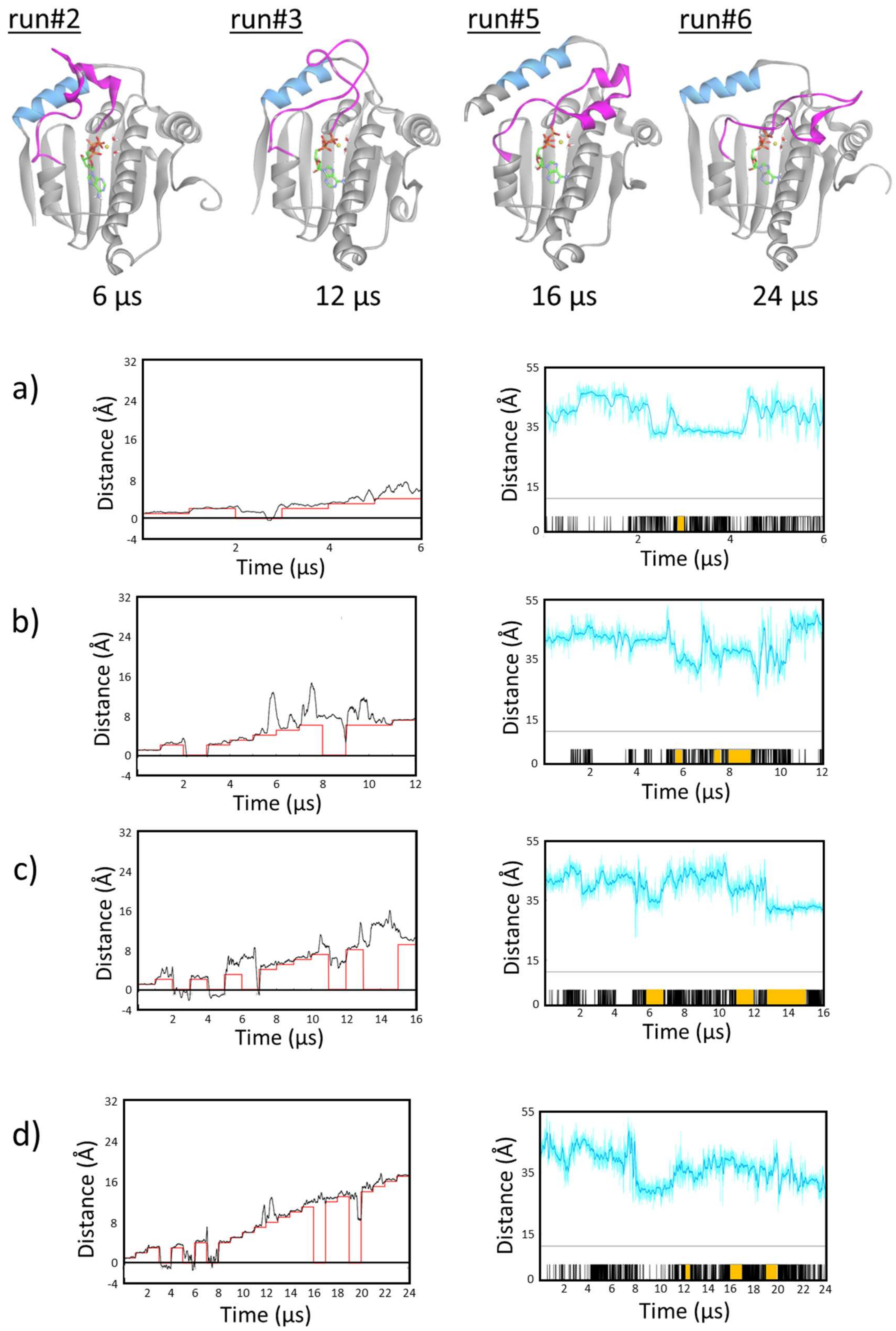


**Figure S2.** Failed flopping-down simulation of the NTD structure with upconformation: runs #2, #3, #5, and #6. Final structures of the flopping-down simulations are shown at the top. a–d) In the graphs on the left, the mean trajectories of the four restrained distances (black lines) and the distance restraints (red lines) are shown. In the graphs of the right, the trajectory of the S51-A110 distance (light-blue lines) is shown. Trajectory in light-green shows the "formation of the down-conformation" period (Section 2.2.2). The "free movement of the lid segment" periods are denoted with black vertical lines and the continuous periods are shown with orange bars (Section 2.2.2). The results are shown in a) runs #2, b) #3, c) #5, and d) #6.

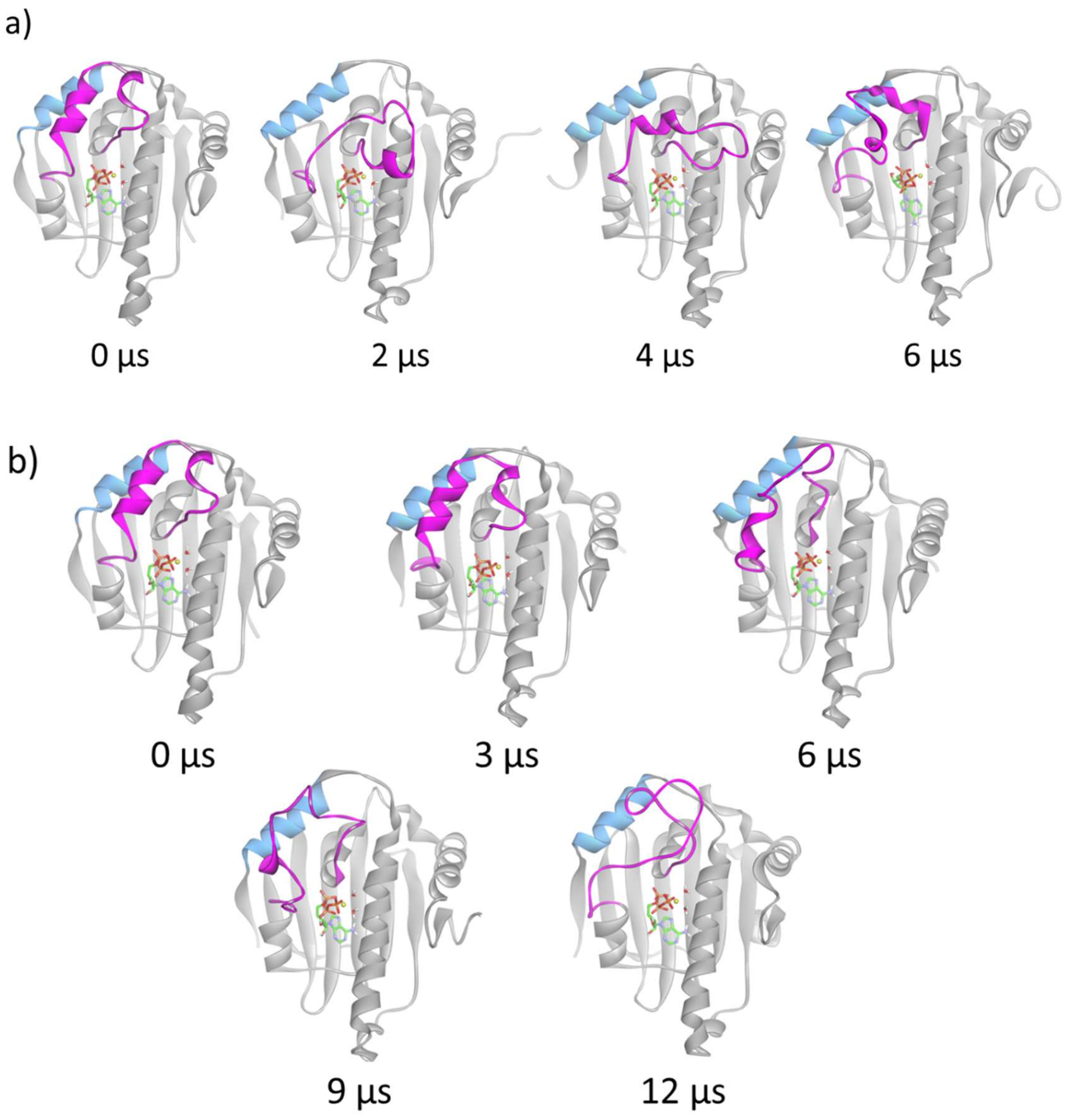


**Figure S3.** Trajectory snapshot structures of the flopping-down simulations, a) runs #2, b) #3, c) #5, and d) #6 from the front view.

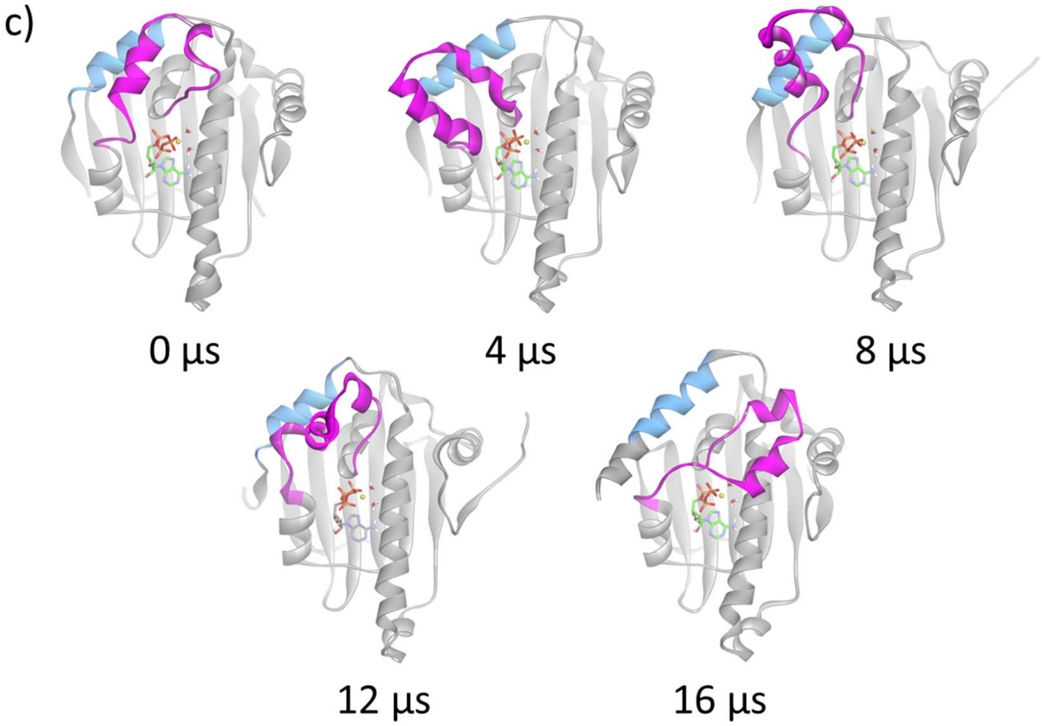


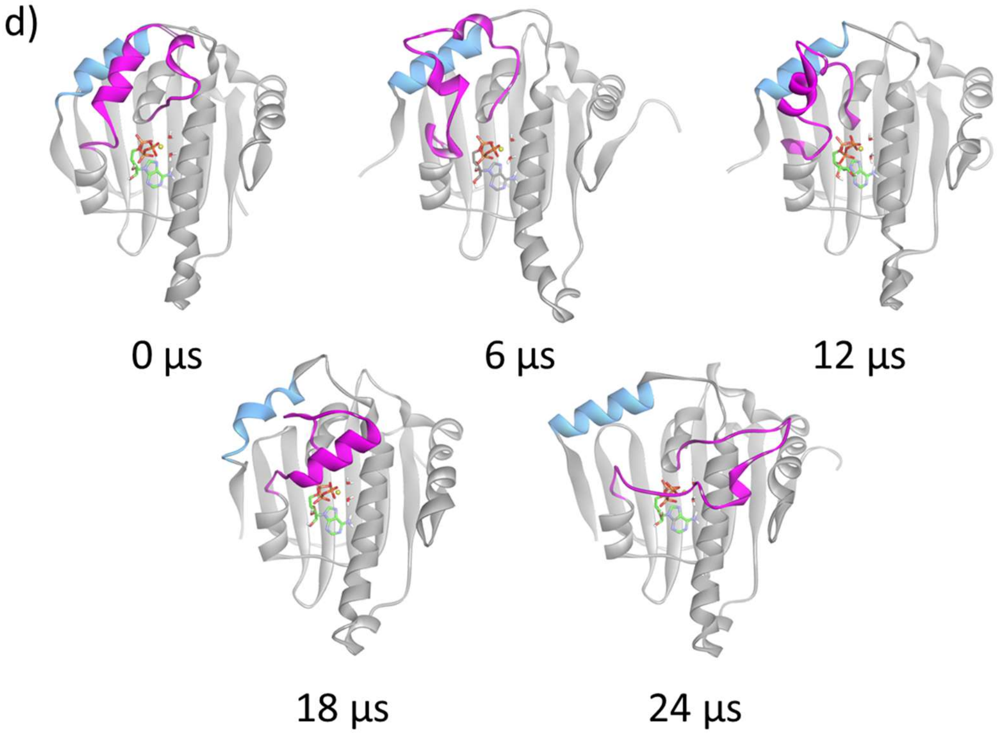


**Figure S3.** Trajectory snapshot structures of the flopping-down simulations, a) runs #2, b) #3, c) #5, and d) #6 from the front view.

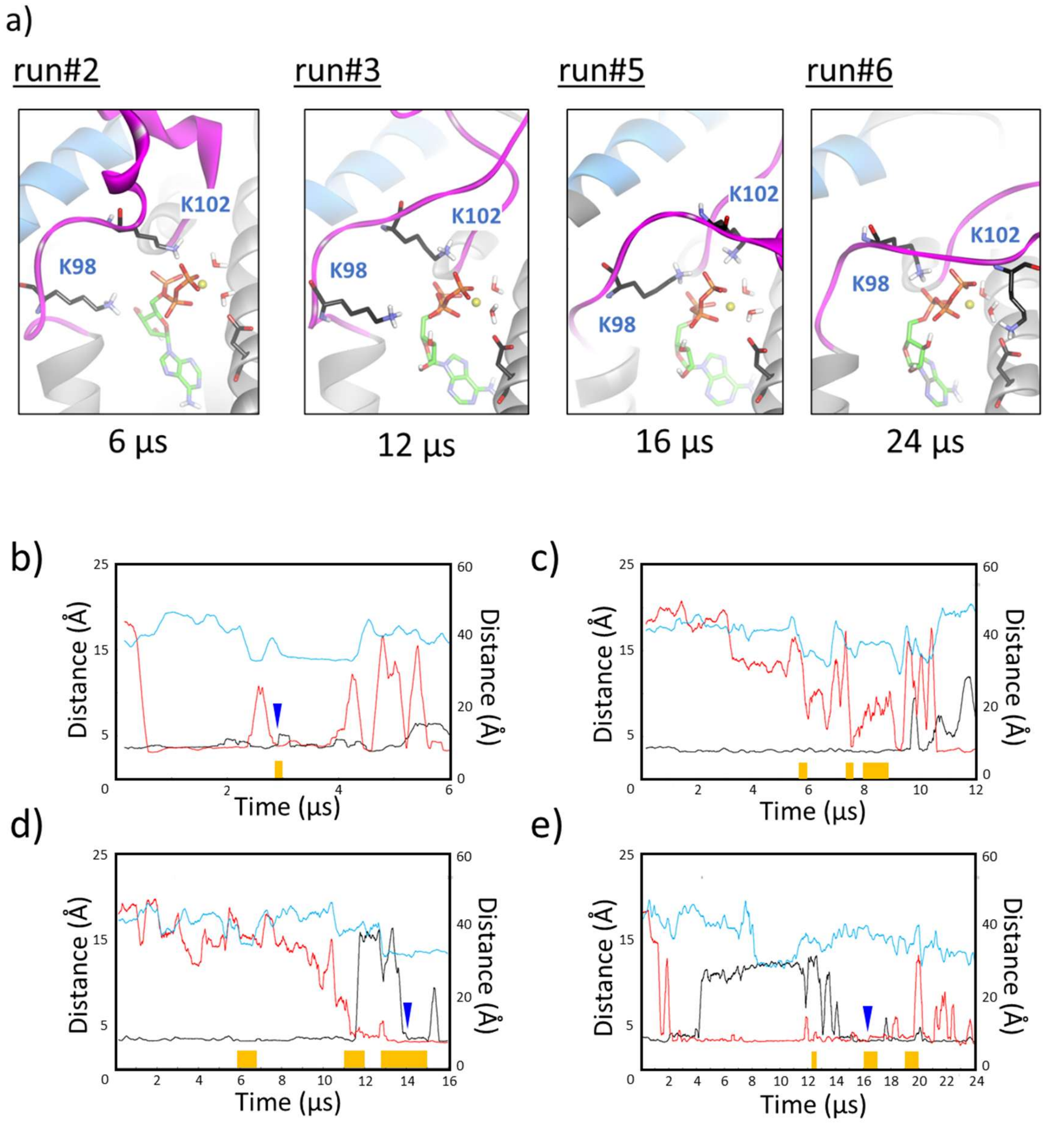


**Figure S4**. Close-up of representative and final structures of the flopping-down simulation, a) runs #2, #3, #5, and #6. b–e) Distance trajectories of K98-ATP phosphate (black lines) and K102-ATP phosphate/D40 (red lies) are shown on the left vertical axis. Distance trajectory of S51-A110 (light-blue lines) is shown on the right vertical axis. Light-green lines and orange bars indicate the periods of the "formation of the down-conformation" and "free movement of the lid segment," respectively, also shown in Figure 2d. Blue wedges in the graphs indicate the starting point of the simultaneous formation of the K98 and K102 interactions under the free movement period of the lid segment.

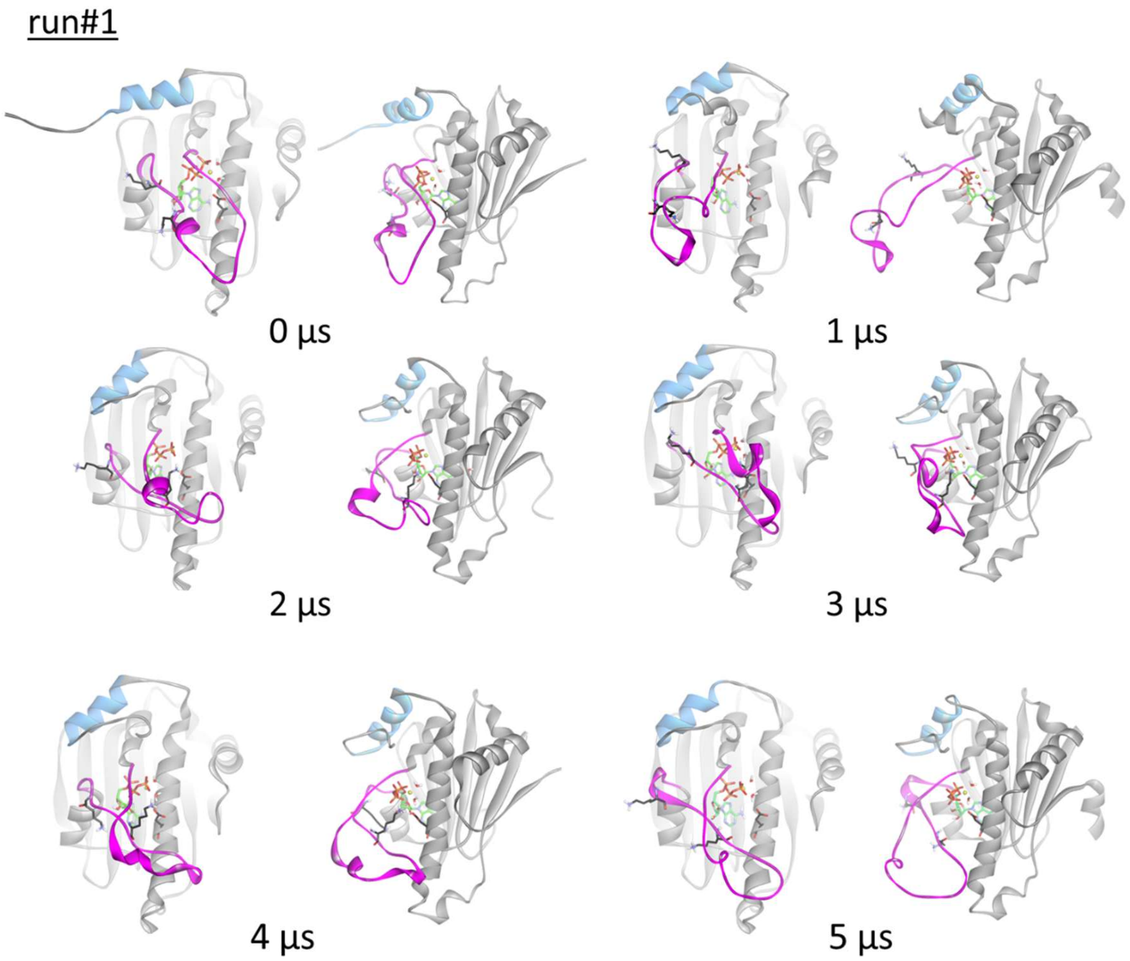


**Figure S5.** Trajectory snapshot structures of the down-conformation simulation, run #1, for the X-ray H1-model from the front and side.

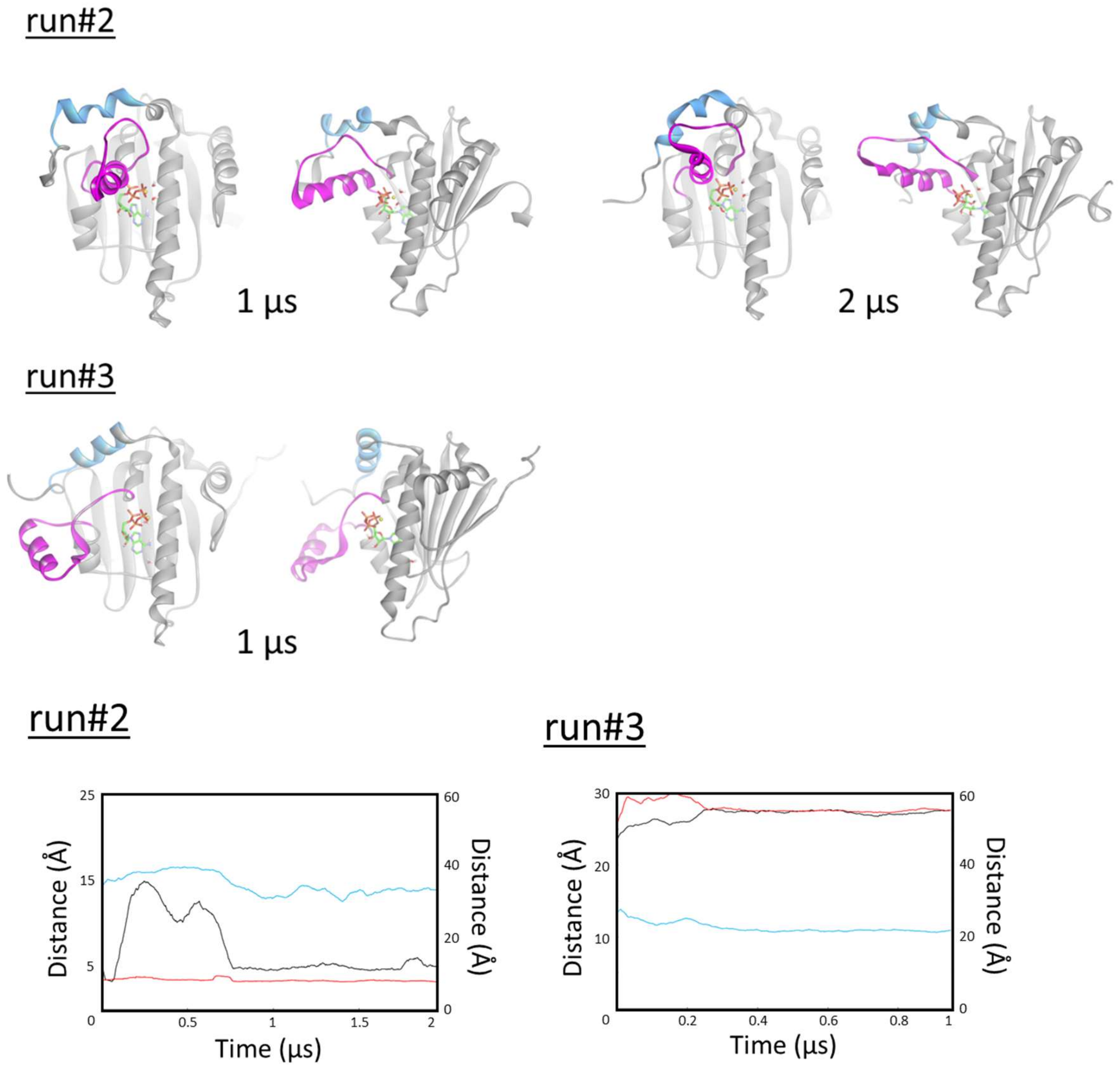


**Figure S6.** Trajectory snapshot structures of the down-conformation simulation, runs #2 and #3, for the X-ray H1-model from the front and side. Distance trajectories of K98-ATP phosphate (black lines) and K102-ATP phosphate/D40 (red lies) are shown on the left vertical axis. Distance trajectory of S51-A110 (light-blue lines) is shown on the right vertical axis.

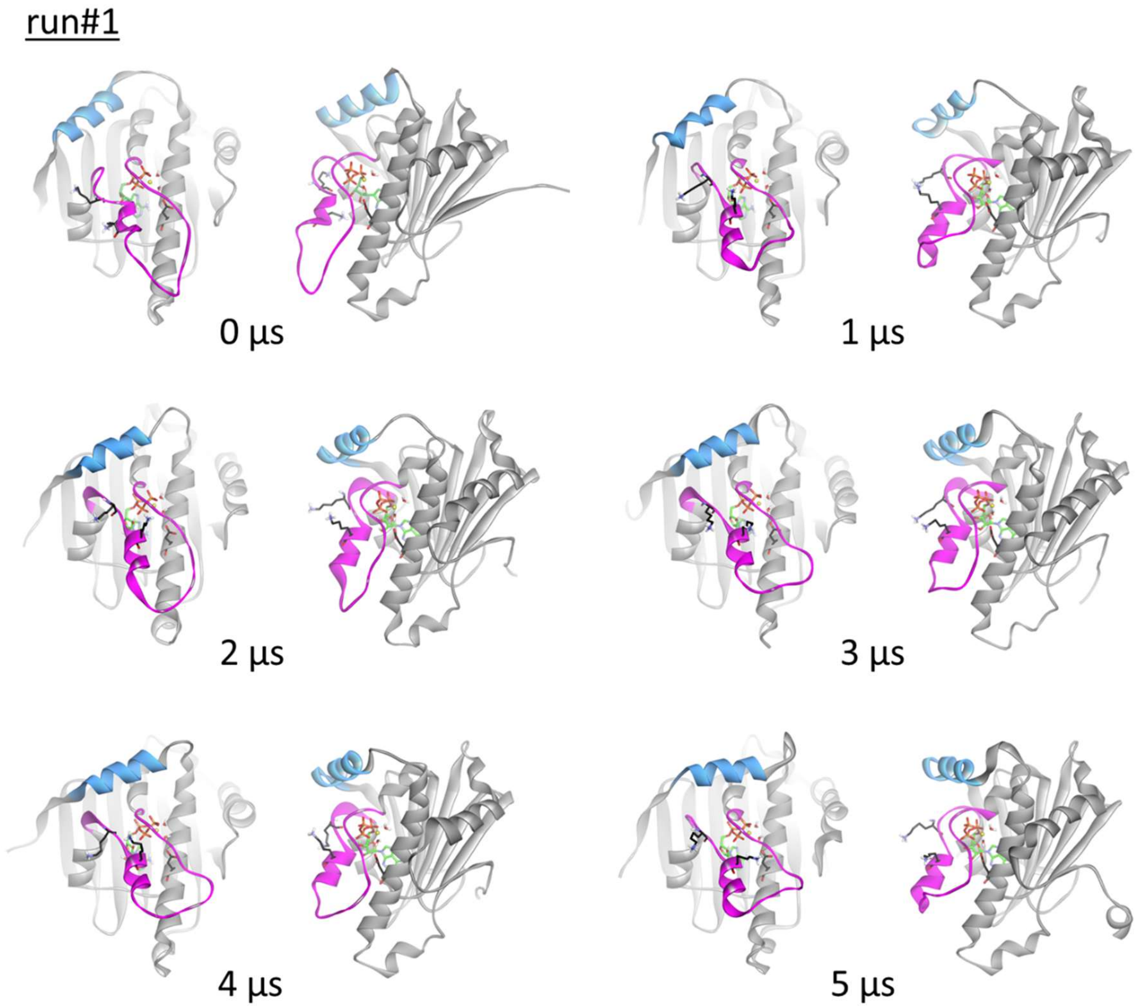


**Figure S7.** Trajectory snapshot structures of the down-conformation simulation, runs #1 and #2, for the chimera H1-structure from the front and side.

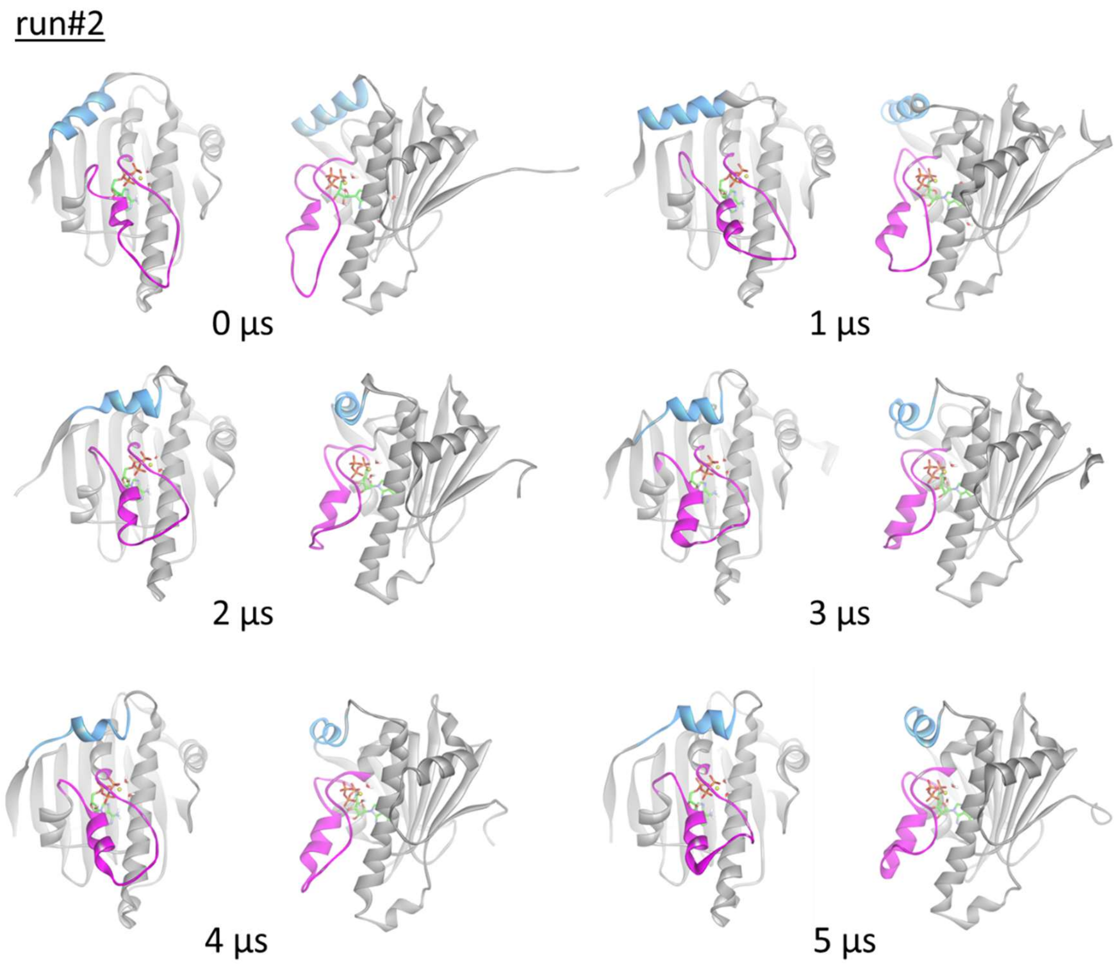


**Figure S7.** Trajectory snapshot structures of the down-conformation simulation, runs #1 and #2, for the chimera H1-structure from the front and side.

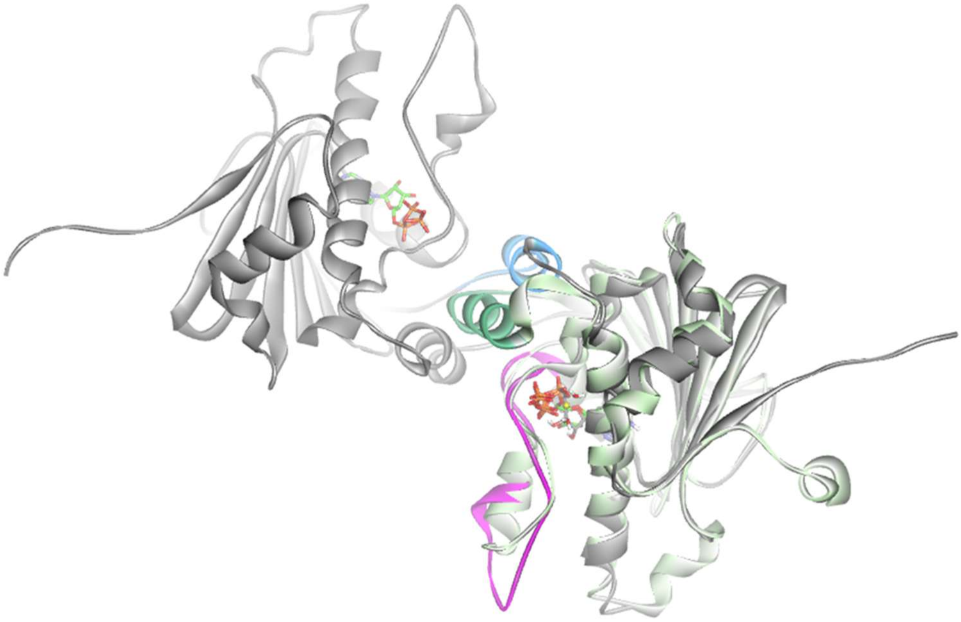

**Figure S8.** Superimposition of the chimeric H1-model. Comparison of the final structures (5 μs) between runs #1 (gray) and #2 (orange). H1 segments are colored blue and gray for the X-ray NTD structure and in green for the chimera H1-model.

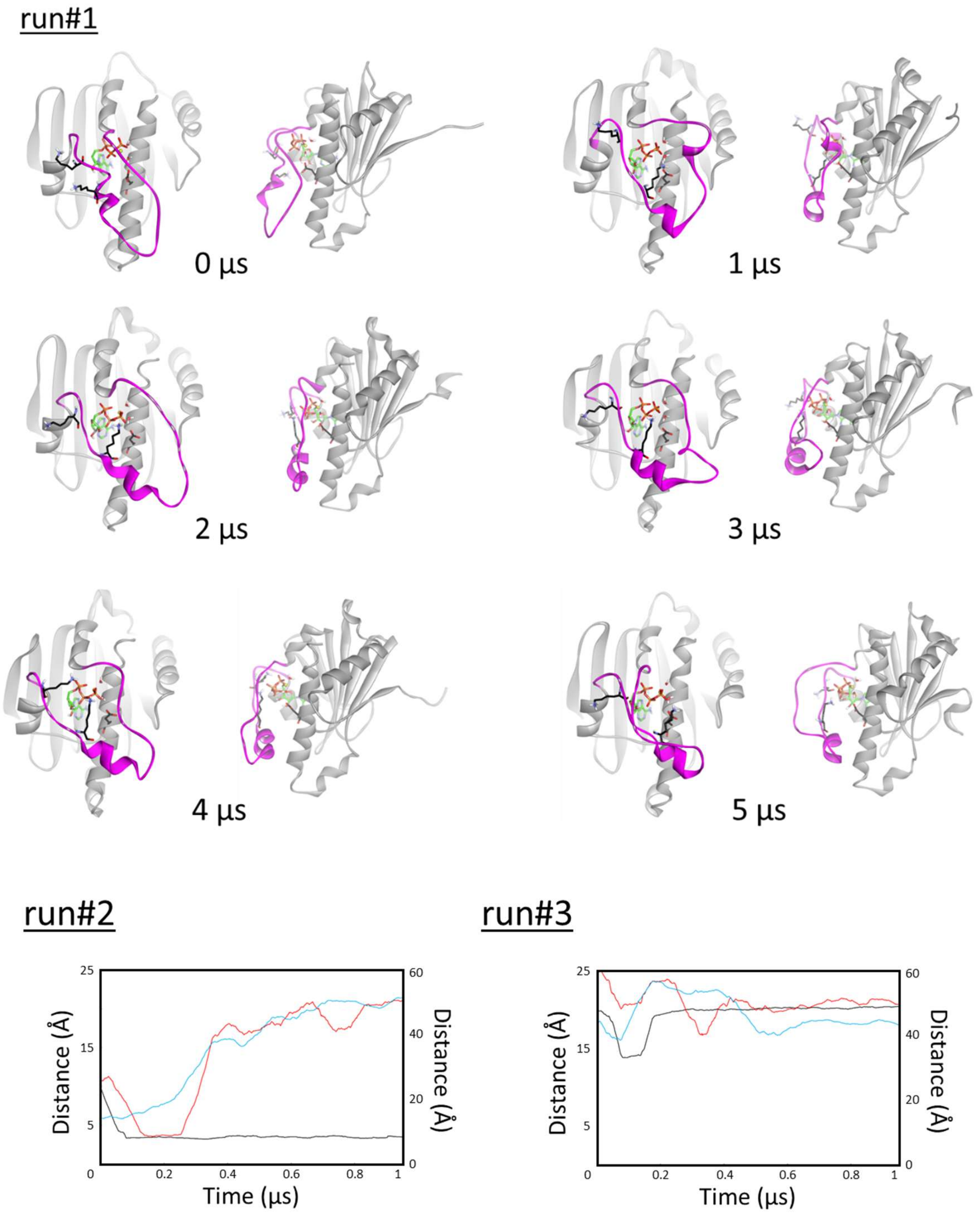


**Figure S9.** Trajectory snapshot structures of the down-conformation simulation, run#1, for the truncated H1-model from the front and side. In the graphs, distance trajectories of K98-ATP phosphate (black lines) and K102-ATP phosphate/D40 (red lines) are shown on the left vertical axis. Distance trajectory of S51-A110 (light-blue lines) is shown on the right vertical axis.

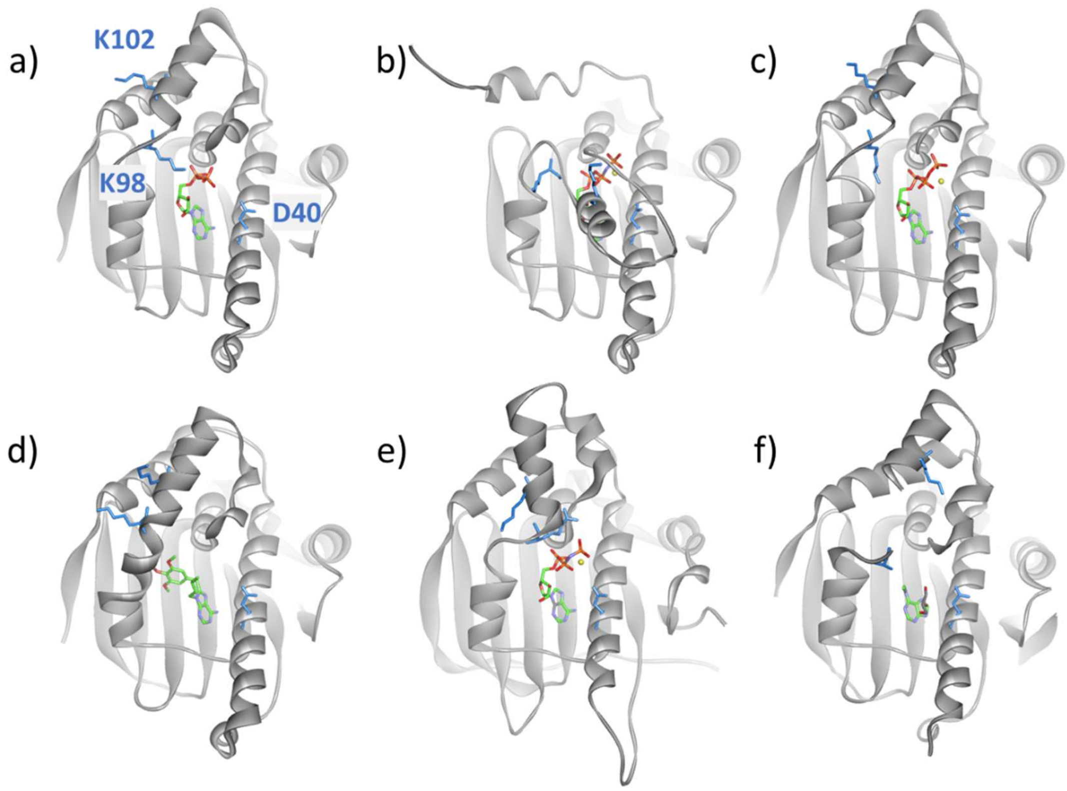


**Figure S10.** Comparison of the corresponding residues of D40, K98 and K102 in Hsp90 homologues that are structurally available. Positions of the residues are as follows: a) D40, K98, and K102 in yeast Hsp82 (1AM1); b) D40, K98, and K102 in yeast Hsc82 (6XLE); c) D54, K112, and K116 in human Hsp90α (3T0Z); d) D49, K107, and K111 in human Hsp90β (6N8Y); e) D122, R177, and K181 in human TRAP1 (5F5R); and f) D110, K168, and K177 in human Grp94 (7ULL). A part of lid segment and side-chain of K168 in human Grp94 are missing because of disorder.

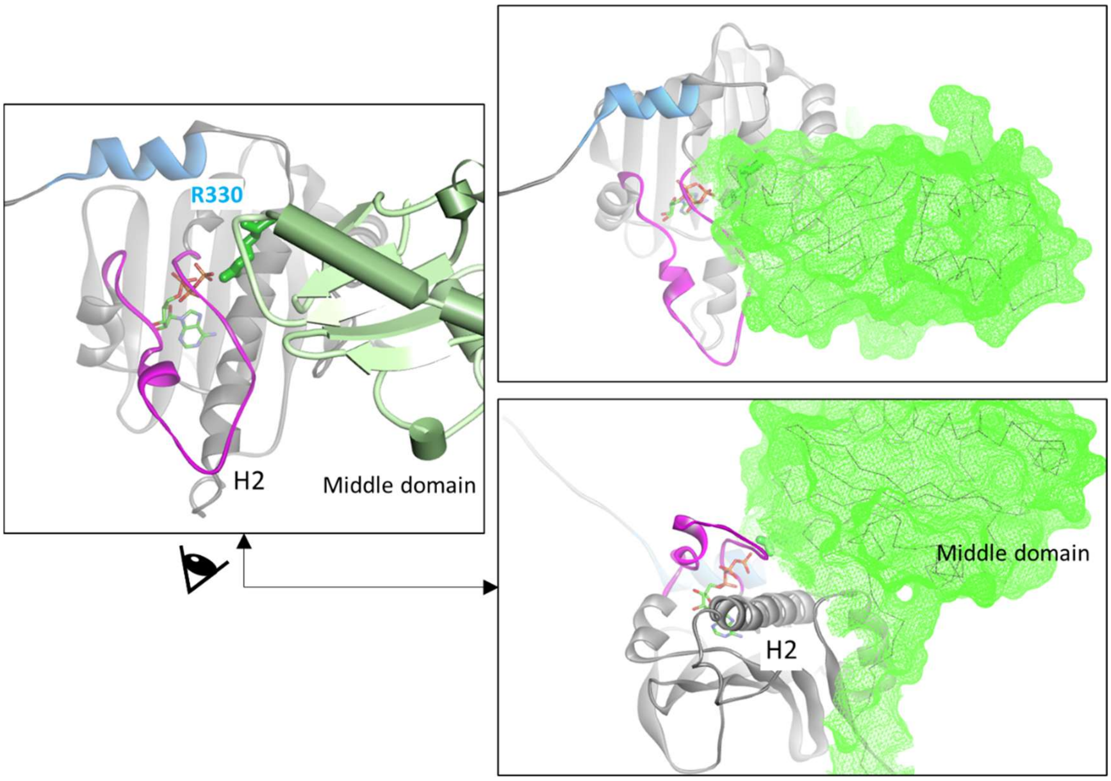


**Figure S11.** X-ray structure of the NTD-middle domain complex in the dimeric Hsp90 (2CG9). Only the monomer of Hsp90 is shown for clarity. Front views of the NTD-middle domain structure are shown in the left (schematic drawing of the middle domain) and in the top right (mesh drawing of the middle domain). The view from the bottom is shown at the bottom right.
